# A novel prognostic metric resolves the *MYCN* enigma *in silico* and points to a biosynthetic regulatory shift driven by *MYCN* amplification in neuroblastoma

**DOI:** 10.1101/2025.11.13.688307

**Authors:** Matteo Italia, Melody Parker, Rory Deignan, Fabio Dercole, Dawn Walker, Kenneth Y. Wertheim

## Abstract

Neuroblastoma (NB) is the most common extra-cranial solid tumour in children. Although *MYCN* amplification is typically indicative of a poor prognosis, the expression level of *MYCN* has a non-monotonic relationship with the clinical outcome. This paper proposes an explanation for this phenomenon, which is called the *MYCN* enigma in the literature. The p53/MYCN metric, which is a measure of the p53 protein level relative to the MYCN protein level in a NB tumour, is the key concept. Our hypothesis is that the metric has a positive relationship with the outcome of a patient. It is presented as a binary classification system, where the prognosis is favourable when p53/MYCN***>***3.5. The mathematical model presented in this paper describes the dynamics between MYCN, p53, ARF, and MDM2; their genes; and their mRNA transcripts. Simulations were carried out by solving the model numerically in a series of initial value problems. The results are aligned with a list of clinical and experimental observations. We extracted association rules with the Apriori algorithm and explored the parametric space stochastically. Assuming that MYCN enables biogenesis in NB tumours, our results support the prediction that *MYCN* amplification shifts the balance towards producing MYCN. When a wild-type tumour is deficient in MYCN, stress responses may restore biogenesis in general and preferentially produce MYCN in a negative feedback loop. Another prediction is that *MYCN* - amplified tumours, without treatment, require hard-to-attain biosynthetic rates and stress responses to achieve good outcomes.

## 1 Introduction

Neuroblastoma (NB) is the most common extra-cranial solid tumour in children [1]: the median age of diagnosis is 573 days [2]. Impaired stem and progenitor cells in the neural crest (a transient structure in the human embryo) are responsible for its pathogenesis [3–5]. Due to mutations such as *MYCN* amplification and *ALK* activation [4, 6], they cannot migrate and differentiate into more specific lineages, such as the sympathoadrenal lineage, which generates the sympathetic nervous system. Instead, they become NB cells and collectively, a malignant solid tumour [4, 6]. A patient’s risk group is determined based on multiple criteria, including their stage defined by the International Neuroblastoma Risk Group (INRG) classification system, their age, their primary tumour’s histological type, and additional clinical and molecular parameters [7]. Patients from the low- and intermediate-risk groups are expected to have a five-year overall survival rate above 70 %, but high-risk patients are expected to succumb to the disease or relapse even after multi-modal therapy at a rate above 50 % [8], whereupon survival for more than three years is rare [9, 10]. Emerging strategies such as personalised therapies [11, 12], combination therapies [13], targeted inhibitors [14, 15], and virotherapy [16] aim to improve survival outcomes.

*MYCN* amplification in NB cells is considered a biomarker indicative of a poor prognosis. However, it is not well-defined. In some studies [17, 18], a cell was considered *MYCN*-amplified (MA) when it had 10 or more copies of the gene (measured by FISH assay signals) and the signals surpassed at least three times the number of control signals (PAX3 probe, 2q35). In other studies [19, 20], a four-fold increase was used as the threshold. According to one source [21], this mutation is generally present in around 25 % of all NB cases. It is also a defining characteristic of high-risk cases [22]. As an oncogene, *MYCN* can activate genes involved in metastasis, survival, proliferation, pluripotency, self-renewal, and angiogenesis [21]. It can also switch off genes responsible for differentiation, cell cycle arrest, and immune surveillance [21]. Importantly, [6] observed in their mouse xenografts that *MYCN* amplification and a specific activating mutation of *ALK* were sufficient to drive NB development from neural crest progenitor cells.

Counter-intuitively, the abundance of *MYCN* mRNA in 91 primary NB specimens was found to vary non-monotonically with that of *TrkA* mRNA [23], a favourable prognostic indicator. It should be noted that being a favourable biomarker does not automatically mean *TrkA* mechanistically suppresses neuroblastoma. In other words, the correlation between the two quantities was found to be positive in some cases and negative in the rest. One possibility is that the relationship depends on another molecular species or even the broader cellular context. Out of the 91 specimens, the wild-type (WT) tumours without *MYCN* amplification displayed a much broader range of *TrkA* expression levels than their MA counterparts. Furthermore, the WT tumours with more *MYCN* mRNA also expressed more *TrkA* mRNA (favourable), but the MA tumours expressed less *TrkA* mRNA (unfavourable) than the WT tumours despite expressing even more *MYCN* mRNA. One interpretation is that an intermediate *MYCN* mRNA level leads to a favourable outcome for some unknown reasons. Motivated by these contradictory results, another study designed a gene signature called *MYCN* -157 [24]. It predicts poor outcomes independently of *MYCN* amplification and expression. In the same study, tumours with similar *MYCN* mRNA levels were found to have very different MYCN protein levels. The results reported in these two papers [23, 24] form the basis of the *MYCN* enigma.

This paper proposes a hypothesis to explain this enigma. Here, we define the *MYCN* enigma as the general observation that a NB patient’s clinical outcome does not depend monotonically on their tumour’s *MYCN* expression level or *MYCN* amplification status. We have compiled a list of specific clinical and experimental observations (section S1 in the supplementary file). A satisfactory explanation must be consistent with them. Our hypothesis is that the p53 protein level in a NB tumour, relative to the MYCN protein level therein, decides whether it is likely to progress or regress. Henceforth, we will simply call this metric p53/MYCN. This hypothesis is based on the known fact that MYCN (protein) activates *p53* [25] and on the consensus that *p53* is a tumour suppressor gene with diverse downstream effects, including DNA repair [26], cell cycle arrest [27–29], and apoptosis [30, 31]. Although *MYCN* (gene) is typically considered an oncogene [21], it activates a tumour suppressor gene (*p53*) too, suggesting a non-monotonic relationship mirroring the *MYCN* enigma. Intuitively, this tension could be the mechanism behind the phenomenon. If our hypothesis was true, a NB with low p53/MYCN would have a poor clinical outcome. For this hypothesis to be consistent with [23]’s dataset, p53/MYCN should follow a concave function of *MYCN*’s mRNA level in a mixed collection of WT and MA NBs.

Although we found numerous modelling studies involving MDM2 and p53 in the literature [32–38], and even some investigating their interactions with ARF and other interaction partners of p53 (such as ATM and WIP1) [39–43], our literature review led to the conclusion that a novel mathematical model coupling MYCN to the ARF/MDM2/p53 axis was needed to explore our hypothesis. We formulated eight non-linear ordinary differential equations (ODEs) to model the dynamics between MYCN, p53, ARF, and MDM2, as well as their corresponding genes and mRNA transcripts. The model describes a stream of genetic information flowing through a molecular network connecting MYCN to the ARF/MDM2/p53 axis, which is highly relevant to NB research [44]. It is also a suitable context for our task. *TrkA* and its associated products are not in the model because they do not interact with the network directly. There is also no evidence that MYCN activates *TrkA* like it activates *p53*, so the p53/MYCN metric is mechanistically more relevant, making it a more direct proxy for a favourable prognosis than the mRNA level of *TrkA*. After parameterising the model, a series of computer simulations were carried out. Based on the results, a binary NB classification system was established: the prognosis is favourable when p53/MYCN*>*3.5. It predicts whether a patient will have a good or poor outcome based on their p53/MYCN metric. The results are consistent with most of the clinical and experimental observations listed above too. The Apriori algorithm [45] and an exploratory data analysis helped us infer plausible mechanisms underlying the hypothesis.

## 2 Methodology

We will begin this section by presenting a model comprising eight non-linear ODEs describing a gene regulatory network formed by *MYCN*, *ARF*, *MDM2*, *p53*, as well as their mRNA transcripts and proteins. After that and a summary of the computer simulations we performed, we will describe in detail the procedures we followed to establish the NB classification system based on our hypothesis and simulation results. We will explain how we extracted further insights from the simulation results with the Apriori algorithm and exploratory data analysis techniques too.

### 2.1 Model Description

The mathematical model comprises four pairs of master equations. Each pair is based on the central dogma of molecular biology, which states that genetic information flows from a cell’s genome to its transcriptome to its proteome [46]. The first step of information transfer is called transcription, while the second is called translation. The master equations are presented and explained in subsection S2.1 in the supplementary file. In this paper, gene and mRNA names are italicised by default, but protein names are not. An exception is that a variable or parameter is always italicised.

The first four equations below govern transcription and mRNA degradation, which can be derived from equation S1. The four equations after them govern translation and protein degradation, which can be derived from equation S2. Transcription, mRNA degradation, translation, and protein degradation are the four fundamental processes in this model.

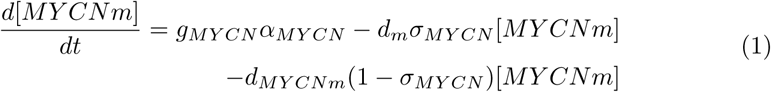

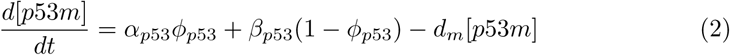

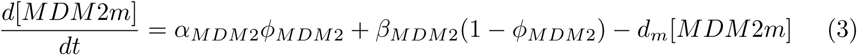

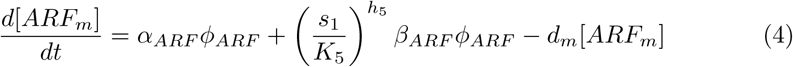

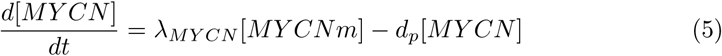

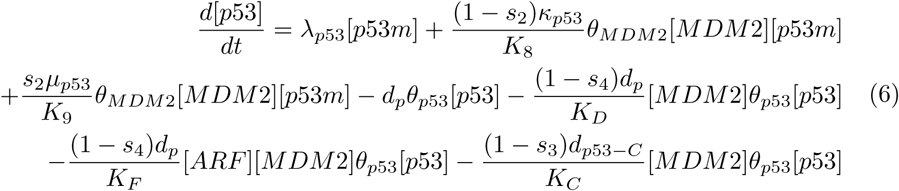

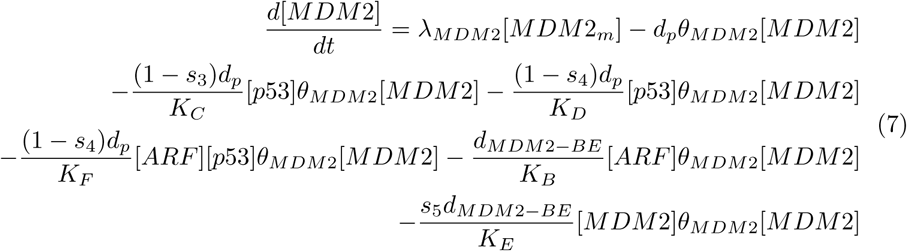

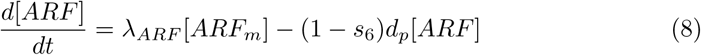

In the following subsections, we will provide the biological basis of the model. Briefly, the equations describe how environmental stressors influence the molecular network by activating or suppressing the protein complexation events illustrated in figure 1, the protein-DNA interactions illustrated in figure 2, and the protein-mRNA interactions illustrated in figure 3, as well as the downstream effects on the four fundamental processes. For example, oncogenic stresses activate *c-Myc* to inhibit ARF degradation [47–49] (edge 1.i, figure 1). In this model, this stressor is denoted by *s*_6_. The detailed modelling procedures are explained in subsections S2.2, S2.3, S2.4, and S2.5 in the supplementary file. For an exposition of modelling gene regulatory networks, chapter 7 of [50] is recommended. Tables 1 and 2 summarise the model’s parameters, including their values, units, and biological descriptions. For a detailed account justifying the chosen values, please refer to subsection S2.6 in the supplementary file.

**Fig. 1:**
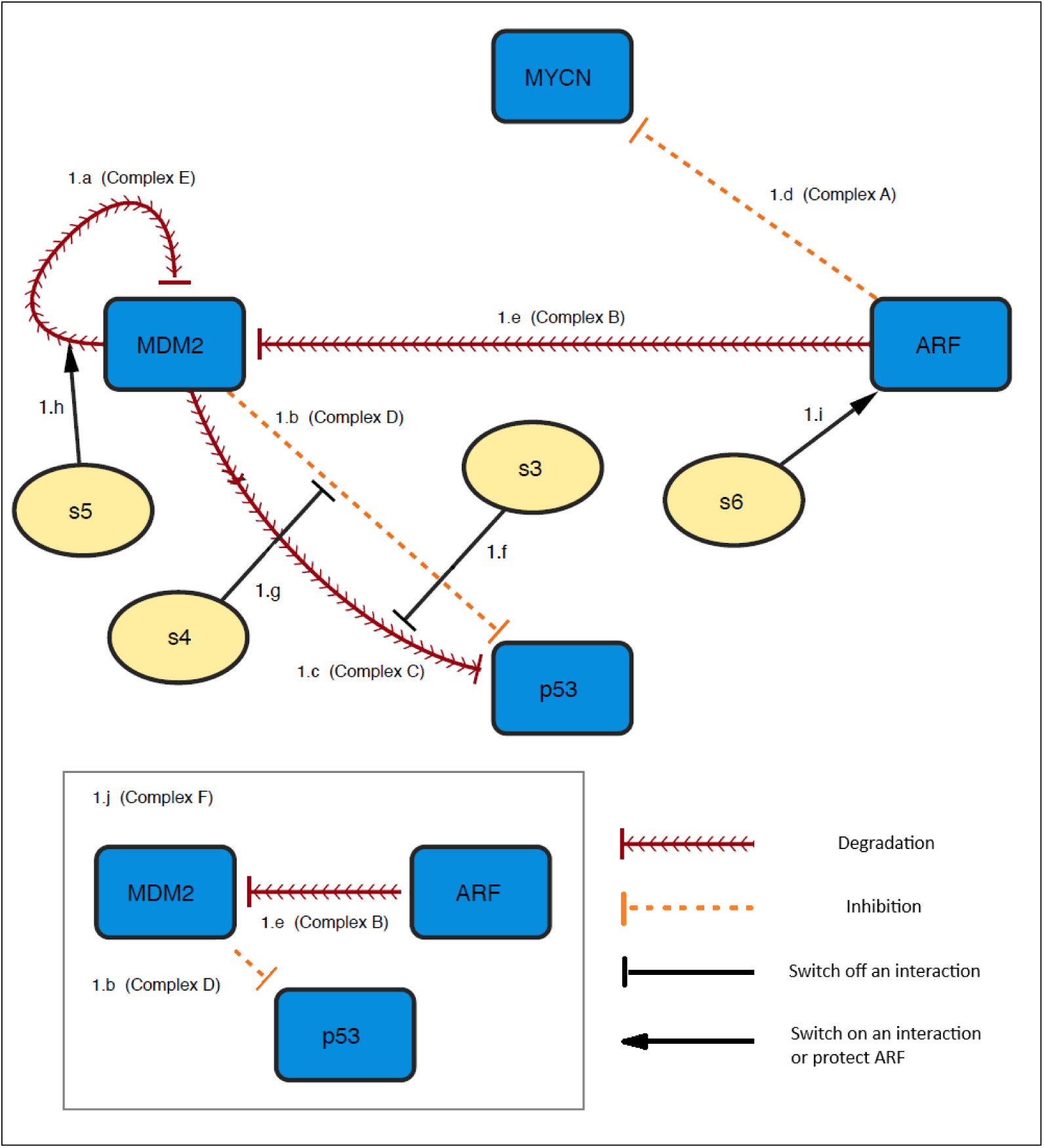
Protein complexation events. (a) MYCN protein’s ability to transcribe *p53* and *MDM2* is downregulated (equations 2 and 3) when it forms complex A with ARF. (b) MDM2 protein is degraded by itself in Complex E and by ARF in complex B and complex F (equation 7). The stressor *s*_5_ promotes complex E formation. (c) p53 protein’s ability to transcribe *p53* and *MDM2* is downregulated (equations 2 and 3) by MDM2 in complex D and complex F. It is also degraded by MDM2 in complex C (equation 6). The stressors *s*_3_ and *s*_4_ prevent complex C and complex D from forming respectively. (d) The stressor *s*_6_ reduces ARF protein’s degradation rate (equation 8).

**Fig. 2:**
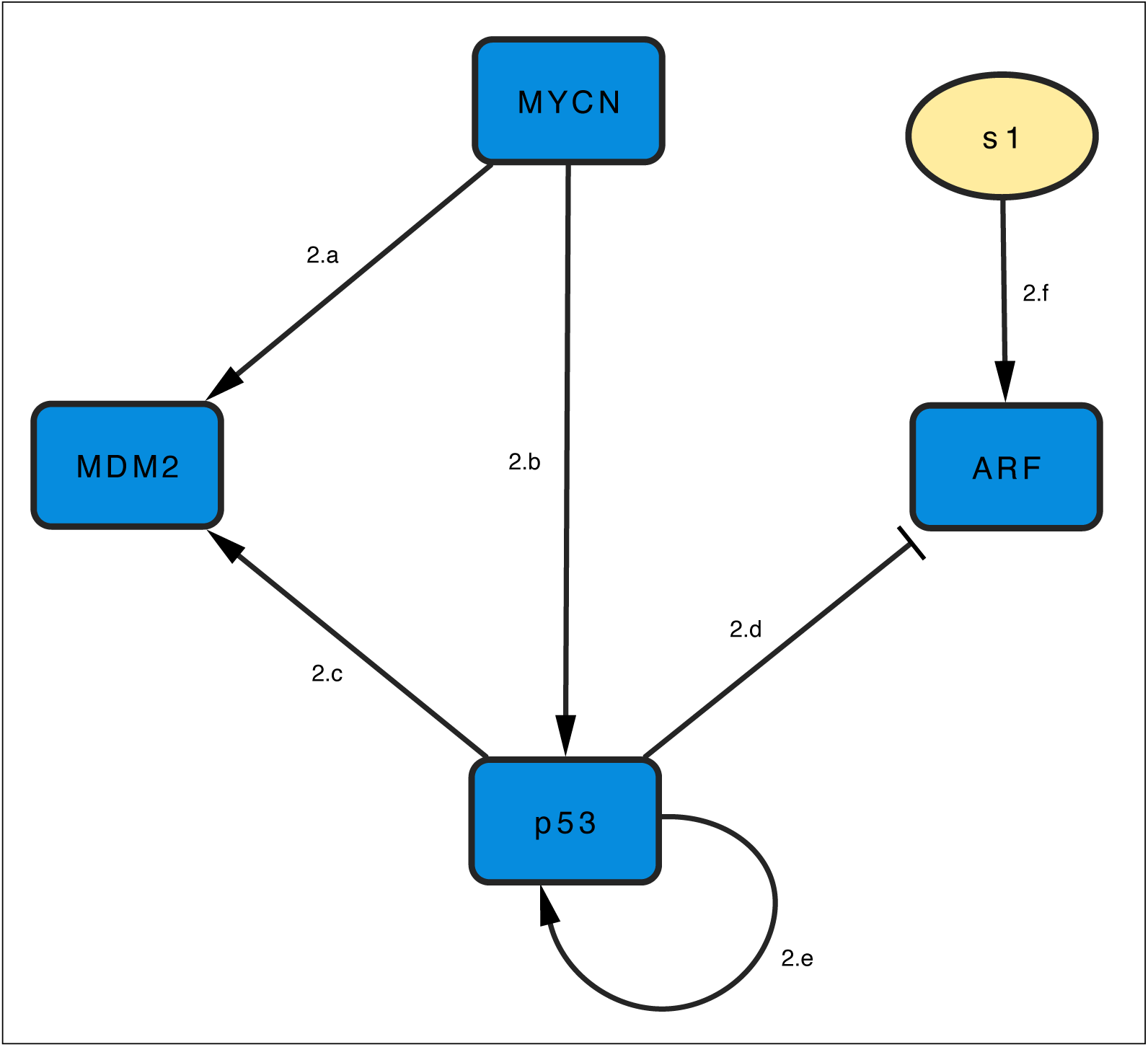
Interactions between proteins and DNA promoter regions. (a) The promoter region of *MYCN* is not regulated by these four proteins (equation 1). (b) The promoter region of *MDM2* is activated (more transcribed) by MYCN and p53 (equation 3). (c) The promoter region of *p53* is activated (more transcribed) by MYCN and p53 (equation 2). (d) The promoter region of *ARF* is inhibited (less transcribed) when occupied by p53 and activated (more transcribed) when occupied by the stressor *s*_1_ (equation 4).

**Fig. 3:**
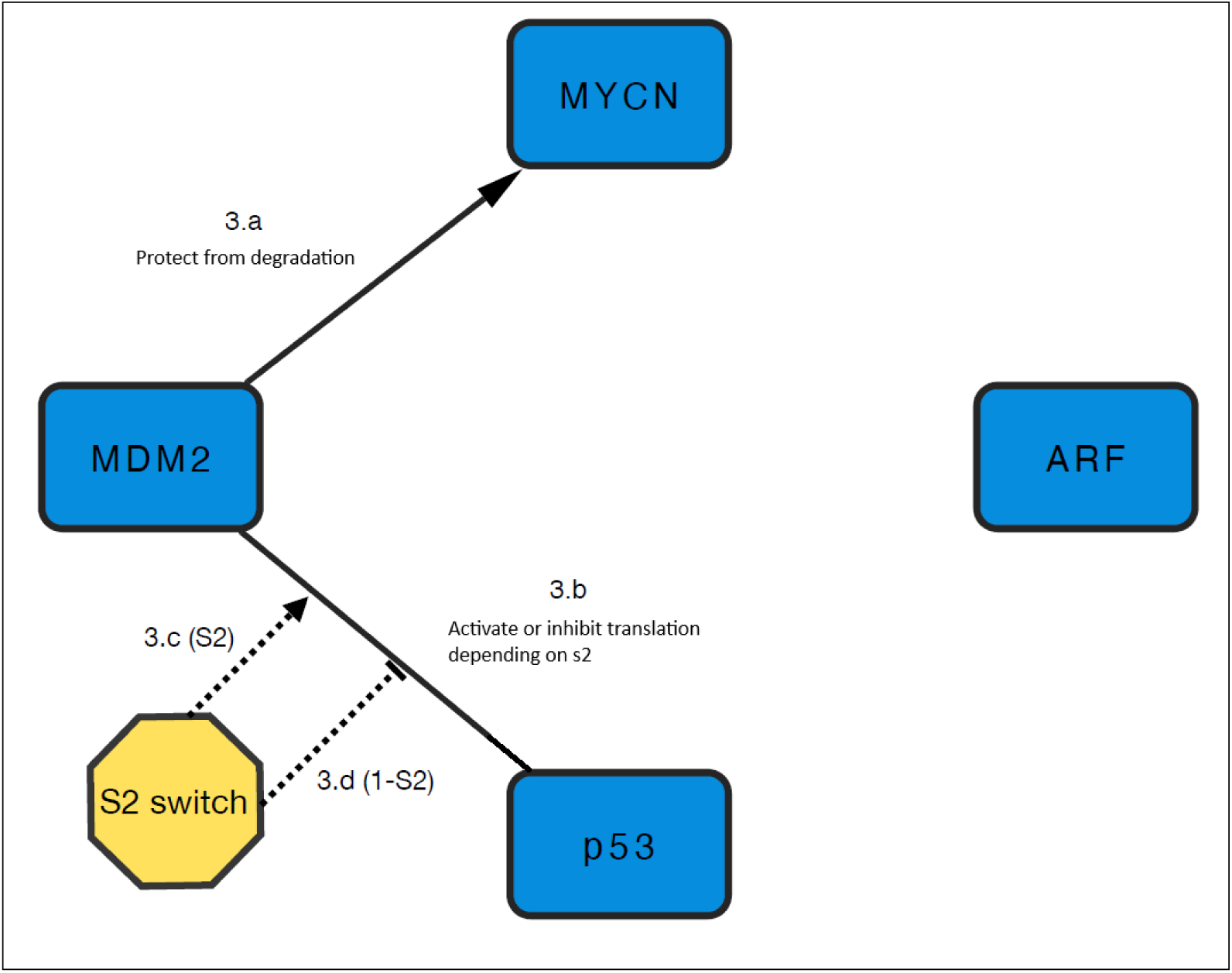
Interactions between proteins and mRNA transcripts. (a) MDM2 binds to and reduces the degradation rate of *MYCN* transcripts (equation 1). (b) The transcripts of *MDM2* are not regulated by these four proteins (equation 3). (c) The rate at which *p53* transcripts are translated is dependent on MDM2, which raises the rate when the stressor *s*_2_ is on and lowers it when *s*_2_ is off (equation 6). (d) The transcripts of *ARF* are not regulated by these four proteins (equation 4).

**Table 1:** Model parameters controlling the four fundamental processes in our model. DR denotes ‘degradation rate’. A denotes ‘assumed’.

| Parameter | Value | Units | Description | Source |
| --- | --- | --- | --- | --- |
| $d_m$ | 0.06 | $\text{h}^{-1}$ | Natural mRNA degradation rate. | [51] |
| $d_{MYCNm}$ | 0.006 | $\text{h}^{-1}$ | *MYCN* mRNA DR when bound to MDM2. | A |
| $d_p$ | 0.05 | $\text{h}^{-1}$ | Natural protein degradation rate. | [51] |
| $d_{p53-C}$ | 0.5 | $\text{h}^{-1}$ | p53 protein degradation rate in complex C. | A |
| $d_{MDM2-BE}$ | 0.5 | $\text{h}^{-1}$ | MDM2 protein DR in complexes B and E. | A |
| $g_{MYCN}$ | 1 or 2 | - | *MYCN* copy number. | A |
| $\alpha_{MYCN}$ | 0.088 | $\text{nM h}^{-1}$ | Natural *MYCN* transcription rate. | [52] |
| $\alpha_{p53}$ | $0.1\beta_{p53}$ | $\text{nM h}^{-1}$ | Natural *p53* transcription rate. | A |
| $\beta_{p53}$ | 0.705 | $\text{nM h}^{-1}$ | Enhanced *p53* transcription rate. | [52] |
| $\alpha_{MDM2}$ | $0.1\beta_{MDM2}$ | $\text{nM h}^{-1}$ | Natural *MDM2* transcription rate. | A |
| $\beta_{MDM2}$ | 1.072 | $\text{nM h}^{-1}$ | Enhanced *MDM2* transcription rate. | [52] |
| $\alpha_{ARF}$ | $0.1\beta_{ARF}$ | $\text{nM h}^{-1}$ | Natural *ARF* transcription rate. | A |
| $\beta_{ARF}$ | 5.034 | $\text{nM h}^{-1}$ | Enhanced *ARF* transcription rate. | [52] |
| $\lambda_{MYCN}$ | 36.23 | $\text{h}^{-1}$ | Natural *MYCN* translation rate. | [53] |
| $\mu_{p53}$ | 2319.68 | $\text{h}^{-1}$ | Enhanced p53 translation rate. | [53] |
| $\lambda_{p53}$ | 231.97 | $\text{h}^{-1}$ | Natural p53 translation rate. | A |
| $\kappa_{p53}$ | 23.2 | $\text{h}^{-1}$ | Reduced p53 translation rate. | A |
| $\lambda_{MDM2}$ | 356.61 | $\text{h}^{-1}$ | Natural MDM2 translation rate. | [53] |
| $\lambda_{ARF}$ | 1603.31 | $\text{h}^{-1}$ | Natural ARF translation rate. | [53] |

**Table 2:** Equilibrium dissociation constants (DC) and Hill coefficient. P denotes ‘parameter’. S denotes ‘source’. A denotes ‘assumed’.

| P | Value | Units | Description | S |
| --- | --- | --- | --- | --- |
| $K_1$ | 20 | nM | DC for MYCN bound to *p53* DNA. | [54] |
| $K_2$ | 121 | nM | DC for p53 bound to *p53* DNA. | [54] |
| $K_3$ | 20 | nM | DC for MYCN bound to *MDM2* DNA. | [54] |
| $K_4$ | 121 | nM | DC for p53 bound to *MDM2* DNA. | [55] |
| $K_5$ | 1 | - | DC for stress-dependent *ARF* transcriptional activation. | A |
| $h_5$ | 1 | - | Hill coefficient for stress activating *ARF* transcription. | A |
| $K_6$ | 121 | nM | DC for p53 bound to *ARF* DNA. | [55] |
| $K_7$ | 200 | nM | DC for MDM2 bound to *MYCN* mRNA. | [56] |
| $K_8$ | 100 | nM | DC for when MDM2 acts as an inhibitor. | [57] |
| $K_9$ | 200 | nM | DC for MDM2 bound to and activating *p53* mRNA. | [56] |
| $K_A$ | 100 | nM | DC for the MYCN-ARF complex A. | A |
| $K_B$ | 100 | nM | DC for the ARF-MDM2 complex B. | A |
| $K_C$ | 77 | nM | DC for the p53-MDM2 complex C. | [58] |
| $K_D$ | 77 | nM | DC for the p53-MDM2 complex D. | [58] |
| $K_E$ | 300 | nM | Maximum DC for the MDM2-MDM2 complex E. | [59] |
| $K_F$ | $K_B K_D$ | nM | DC for the p53-MDM2-ARF complex F. | A |

#### 2.1.1 Environmental stressors

A NB cell responds to a multitude of environmental stressors, which elicit intracellular molecular changes. For example, they may inhibit protein complexation events selectively. They can be thought of as the model’s inputs, but for the sake of simplicity, they are classified as parameters in this paper. There are six stressors, which are denoted by the parameters *s*_1_, *s*_2_, *s*_3_, *s*_4_, *s*_5_, and *s*_6_. Their effects on protein-protein, protein-DNA, and protein-mRNA interactions are illustrated in figures 1, 2, and 3, respectively. They are dimensionless and lie on the interval [0, 1]. As a stressor increases from zero to one, the cell transitions from a stress-free environment to a maximum stress environment. Here are the intracellular changes induced by them and the molecular mechanisms that allow them to do so.

- *s*_1_ is linked to an increased *ARF* transcription rate (edge 2.f, figure 2). This parameter represents oncogenic stresses that activate *ARF* transcription through E2F-1 [60–62] and Ras [63, 64], as well as oxidative and heat shock stresses [65].
- *s*_2_ is linked to an increased *p53* translation rate through MDM2 through two mechanisms (collectively represented by edge 3.c, figure 3). *p53* translation is inhibited or induced by MDM2 depending on the cellular context. MDM2 naturally inhibits the translation activator L26 (scenario B, figure 4 or edge 3.d, figure 3), which would otherwise activate *p53* translation (scenario A, figure 4). In the first mechanism (scenario D, figure 4), stresses in general prevent this negative action of MDM2 on *p53* translation [66–68]. In the second mechanism (also scenario D, figure 4), specific stresses such as DNA damage induce MDM2 to bind to *p53* mRNA transcripts. This process activates translation [69, 70]. In other words, these specific stresses promote MDM2’s positive action on *p53* translation.
- *s*_3_ is linked to a reduced p53 degradation rate (edge 1.f, figure 1). In a stressed cell, both MDM2 and p53 are phosphorylated and acetylated to prevent complexation (complex C in subsection 2.1.2), which normally induces p53 degradation [71, 72]. For example, ATM phosphorylates MDM2 in response to DNA damage induced by radiomimetic chemicals and ionising radiation, but not UV radiation [73]. ATM is a part of *s*_3_.
- *s*_4_ prevents MDM2 from forming a complex with p53 (complex D and complex F in subsection 2.1.2) through phosphorylation and acetylation [72] (edge 1.g, figure 1). In the complexes, p53’s transactivation function is blocked, so *s*_4_ enhances this function [72].
- *s*_5_ is linked to an increased MDM2 degradation rate (edge 1.h, figure 1). Cellular stresses signal through HAUSP to MDM2 [74], making MDM2 switch functionally from degrading p53 to dimerising with and degrading itself (complex E in subsection 2.1.2) [75].
- *s*_6_ is linked to a reduced ARF degradation rate (edge 1.i, figure 1). Oncogenic stresses activate *c-Myc*, which inhibits ARF’s ULF-dependent degradation pathway [47–49].

**Fig. 4:**
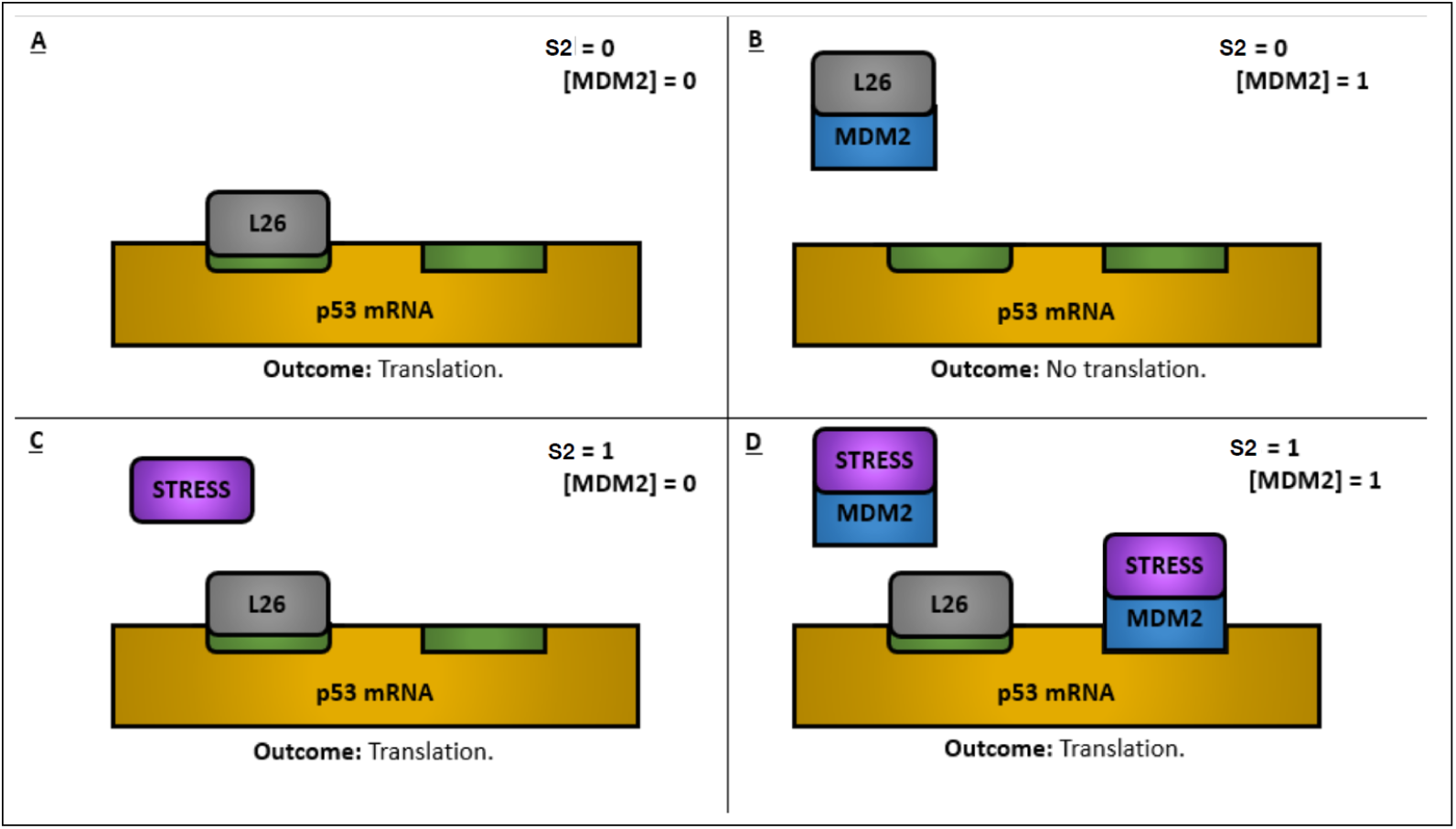
*p53* translation depends on MDM2 and whether the cell is stressed. (A) Naturally, L26 (a ribosomal protein) binds to *p53* mRNA to promote translation. (B) In an unstressed cell, MDM2 binds to L26 to prevent it from promoting *p53* translation. (C) In a stressed cell without MDM2, L26 acts as it does in scenario A to promote translation. (D) In a stressed cell with MDM2, MDM2 does not interact with L26, which can now promote *p53* translation. MDM2 can also bind to *p53* mRNA to promote translation directly. Note that this figure serves to illustrate the biological mechanisms, but in the mathematical model, the functions of L26 are treated in an abstract manner.

In summary, *s*_1_ targets the promoter region of *ARF* ; *s*_2_ targets *p53* mRNA transcripts; *s*_3_, *s*_4_, and *s*_5_ target MDM2 and p53 to influence complexation; and *s*_6_ targets ARF.

#### 2.1.2 Protein complexation

Complexation occurs when two or more proteins bind together to form a larger molecular structure called a complex. Complexation introduces non-linearity to the system because a binding event makes the concentrations of two or more proteins codependent and the resulting complex may exert further effects on the gene regulatory network by modulating the four fundamental processes.

In the list below, we define six protein-protein complexes associated with the gene regulatory network. They are graphically depicted in figure 1. A short description of the binding event(s) is provided for each complex. Note that it is possible for a protein to have multiple binding terminals. For example, complex C and complex D (edges 1.c and 1.b respectively, figure 1) share the same constituents (MDM2 and p53), but they bind at two different terminals in the two complexes. Unsurprisingly, they influence the gene regulatory network differently.

- **Complex A: MYCN and ARF.** In this complex, MYCN is bound to ARF with an equilibrium dissociation constant denoted by *K_A_*. ARF inhibits the transactivation function of MYCN [76], as represented by edge 1.d in figure 1.
- **Complex B: MDM2 and ARF.** In this complex, ARF (either N-terminal or C-terminal) is bound to MDM2 (C-terminal) [77, 78] with an equilibrium dissociation constant denoted by *K_B_*. ARF induces MDM2 degradation, as represented by edge 1.e in figure 1.
- **Complex C: MDM2 and p53.** MDM2 (C-terminal RING finger domain) is bound to p53 (N-terminal) [75, 79, 80] with an equilibrium dissociation constant denoted by *K_C_*. MDM2 induces p53 degradation [81], as represented by edge 1.c in figure 1.
- **Complex D: MDM2 and p53.** MDM2 (N-terminal) is bound to p53 (N-terminal transactivation domain) [72, 82–84] with an equilibrium dissociation constant denoted by *K_D_*. This interaction switches off p53’s transactivation function, as represented by edge 1.b in figure 1. Although p53 is thought to activate and inhibit transcription differently [85, 86], it is assumed in our model that only unbound p53 can get close enough to inhibit transcription. Therefore, this complexation event prevents p53 from influencing transcription in all manners.
- **Complex E: MDM2 and MDM2.** Two copies of MDM2 form this complex with an equilibrium dissociation constant denoted by *K_E_*. In this dimer, MDM2 undergoes self-degradation [75], as represented by edge 1.a in figure 1.
- **Complex F: ARF, MDM2, and p53.** ARF, MDM2 and p53 form a ternary complex. It is assumed that MDM2 is the bridge binding ARF with its C-terminal (as in complex B) and p53 with its N-terminal (as in complex D) [78, 87]. Based on this assumption, the equilibrium dissociation constant is *K_F_* = *K_B_K_D_*. It is also assumed that, as in complex D, this complexation event prevents p53 from influencing transcription in all manners (transactivation and inhibition). How complex F is a combination of complexes B and D is illustrated in inset 1.j in figure 1.

The above complexation events are represented as quasi-equilibrium processes in the model, based on the assumption that protein-protein interactions occur on a much faster timescale than transcription, translation, and degradation. The mathematical details are provided in subsection S2.2 in the supplementary file.

As explained in subsection S2.3 in the supplementary file, these interactions are modelled similarly to protein complexation, but the equilibrium dissociation constants (table 2) are labelled numerically rather than alphabetically.

#### 2.1.3 Interactions between proteins and DNA promoter regions

In addition to protein complexation events, other binding events are non-linear phenomena too. Some of the four proteins can feedback to interact with the genome (DNA) by binding to certain promoter regions to control the relevant transcription rates. Figure 2 summarises how each protein activates or inhibits the promoters of the genes encoding the other three proteins.

- The promoter region of *MYCN* is assumed to be unbound since there is no evidence to the contrary. As a result, there is only one transcription rate for this gene.
- The *p53* gene is transcribed by two transcriptional activators: MYCN [25] (edge 2.b, figure 2) and p53 itself [88] (edge 2.e, figure 2). They bind to non-overlapping sites in the gene’s promoter region [89]. As stated in subsection 2.1.2, MYCN’s transcriptional activity is inhibited by ARF in complex A (edge 1.d, figure 1) and p53’s transcriptional activity is inhibited by MDM2 in complexes D (edge 1.b, figure 1) and F (inset 1.j, figure 1). In complex C (edge 1.c, figure 1), MDM2 induces p53 degradation through the process of ubiquitination. It is fair to assume that p53 cannot transcribe a gene whilst degrading, hence p53 can bind to promoter regions only when it is unbound. Taken together, the *p53* gene is transcribed at a higher rate when its promoter region is occupied by unbound MYCN and/or unbound p53.
- The *MDM2* gene is transcribed by two transcriptional activators: MYCN [90] and p53 [91] (edges 2.a and 2.c respectively, figure 2). Its promoter region has the same binding pattern as that of *p53*, meaning the *MDM2* gene is transcribed at a higher rate when its promoter region is occupied by unbound MYCN and/or unbound p53. However, the binding affinities differ between the two promoter regions.
- The *ARF* gene is down-regulated by p53 [49, 92, 93] and upregulated by a group of proteins activated by the environmental stressor *s*_1_, such as E2F-1, *β*-catenin, and Ras [60, 64, 65], as represented by edges 2.d and 2.f in figure 2 respectively. The gene is transcribed at a natural rate when its promoter region is unoccupied, at a higher rate when it is activated by *s*_1_, and the transcription rate drops to zero when p53 occupies the promoter region.

#### 2.1.4 Interactions between proteins and mRNA transcripts

Figure 3 graphically explains how MDM2 regulates the translation processes producing MYCN and p53. When MDM2 binds to *MYCN* mRNA transcripts, it reduces their degradation rate [94] (edge 3.a, figure 3). It is assumed that when *MYCN* mRNA is bound to MDM2, ribosomal subunits and initiation factors can still bind to it to initiate translation [95, 96]. There is no evidence that the other three mRNA transcripts are protected from or encouraged towards degradation by proteins.

On the other hand, MDM2 affects the rate at which *p53* mRNA transcripts are translated (edge 3.b, figure 3). In an unstressed cell (edge 3.d, figure 3), *p53* translation is indirectly inhibited because MDM2 is free to inhibit L26, which activates *p53* translation [66–68]. In response to *s*_2_ (associated with damaged DNA), L26 is free to promote *p53* translation, which is also directly activated by MDM2 [69, 70]; these positive effects are collectively represented by edge 3.c in figure 3.

The technical details regarding how these interactions are modelled mathematically, including how the effects of L26 are represented abstractly since its concentration is not a state variable, are provided in subsection S2.4 in the supplementary file.

#### 2.1.5 Functional effects of complexation

Our assumptions relating to the effects of complexation on transcription are as follows.

- *MYCN* is transcribed at one rate only.
- *p53* is transcribed at two rates. The natural rate applies when its promoter region is unoccupied. The enhanced rate is achieved when uncomplexed MYCN (edge 2.b, figure 2) and/or uncomplexed p53 (edge 2.e, figure 2) occupies the region.
- *MDM2* gene is transcribed at two rates. The natural rate applies when its promoter region is unoccupied. The enhanced rate is achieved when uncomplexed MYCN (edge 2.a, figure 2) and/or uncomplexed p53 (edge 2.c, figure 2) occupies the region.
- *ARF* is transcribed at three rates. When uncomplexed p53 occupies its promoter region, it is zero (edge 2.d, figure 2). When the promoter is unoccupied, transcription proceeds at the natural rate. When the gene is activated by *s*_1_, an enhanced transcription rate is achieved (edge 2.f, figure 2).

Our assumptions relating to the effects of complexation on mRNA degradation are as follows.

- *MYCN* mRNA transcripts exist in two states, which degrade at a natural rate and a reduced rate respectively. In the uncomplexed form, they degrade naturally. When a transcript is bound by MDM2, it degrades at the reduced rate (edge 3.a, figure 3).
- The other three mRNA transcripts have a fixed degradation rate.

Our assumptions relating to the effects of complexation on translation are as follows.

- Although *MYCN* mRNA exists in two forms (unbound or bound by MDM2), it is assumed that ribosomal subunits and initiation factors can bind to it in both cases to initiate translation [95, 96]. Therefore, it has just one translation rate.
- p53 protein is produced at three rates: natural, enhanced, and reduced. When *p53* mRNA is unbound, translation occurs at the natural rate. When MDM2 binds to it in response to *s*_2_ (associated with damaged DNA), the enhanced rate applies (edge 3.c, figure 3). In an unstressed cell, MDM2’s action leads to a reduced rate of p53 production (edge 3.d, figure 3).
- MDM2 and ARF are produced at the same fixed translation rate.

Finally, our assumptions relating to the effects of complexation on protein degradation are as follows.

- MYCN exists either in its unbound state or in complex A (bound to ARF), as represented by edge 1.d in figure 1. The same natural degradation rate applies in both cases.
- p53 exists in four states: unbound, complex C (bound to MDM2), as represented by edge 1.c in figure 1, complex D (bound to MDM2), as represented by edge 1.b in figure 1, and complex F (bound to ARF and MDM2), as illustrated in inset 1.j in figure 1. It degrades at an enhanced rate in complex C because MDM2 induces p53 degradation. In the other three states, it degrades naturally.
- MDM2 exists in six states: unbound, complex B (bound to ARF), as represented by edge 1.e in figure 1, complex C (bound to p53), as represented by edge 1.c in figure 1, complex D (bound to p53), as represented by edge 1.b in figure 1, complex E (two copies of MDM2), as represented by edge 1.a in figure 1, and complex F (bound to ARF and p53), as illustrated in inset 1.j in figure 1. In complexes B and E, it degrades at an enhanced rate. In the other states, it degrades naturally.
- ARF exists in four states: unbound, complex A (bound to MYCN), as represented by edge 1.d in figure 1, complex B (bound to MDM2), as represented by edge 1.e in figure 1, and complex F (bound to MDM2 and p53), as illustrated in inset 1.j in figure 1. In all four cases, ARF degrades naturally.

How these functional effects are represented mathematically in the model is explained in subsection S2.5 in the supplementary file.

### 2.2 Numerical simulations

We conducted three sets of *in silico* experiments. The configurations and computational details of these experiments can be found in subsection S2.7 in the supplementary file. First, we varied three groups of parameters with and without *MYCN* amplification in a grid search, meaning each parameter was set at predefined values within a range. The results were used to confirm the model’s stable long-term behaviour (equilibrium), which represents the clinical/treatment outcome; understand the outcome’s sensitivity to the three groups of parameters; and establish a NB classification system based on our hypothesis. Second, we focused on the two most sensitive groups of parameters, repeating the grid search without *MYCN* amplification and at five levels of *MYCN* amplification. Third, we managed to reach the conclusions of the first two experiments by exploring the parametric space randomly.

### 2.3 NB classification system based on our hypothesis

In the first experiment, we tested 1259715 parametric combinations by using each to perform two sets of 10 simulations, once with *g_MYCN_* = 1 and once with *g_MYCN_* = 2. In each case, all 20 simulations ended in the same equilibrium state (eight-dimensional fixed point). Furthermore, the 2 · 1259715 = 2519430 fixed points were all found to be stable by perturbing the state vector from and checking that it returned to every fixed point. Then, we applied our hypothesis to the same dataset to create a NB classification system for the purpose of prognosing NB patients.

#### 2.3.1 p53/MYCN: a new prognostic metric

As stated in the introduction, our hypothesis is that the p53 protein level in a NB tumour, relative to the MYCN protein level therein, decides whether it is likely to progress (poor outcome) or regress (good outcome). This hypothesis rests on the fact that *p53* and *MYCN* are, respectively, commonly considered to be a tumour suppressor gene and an oncogene with opposite effects.

We applied the hypothesis to two simulations conducted in the first experiment. They were conducted with the default parameters and inactive stressors (*s_i_* = 0), one with *g_MYCN_* = 1 and the other with *g_MYCN_* = 2. The arithmetic mean of the two equilibrium values of p53/MYCN was calculated and used to define a critical threshold. A patient or simulated equilibrium whose p53/MYCN is above the critical threshold is predicted to have a good outcome. Otherwise, a poor outcome is predicted.

#### 2.3.2 *MYCN* mRNA and MYCN protein levels

We wanted to check if the classification system and by extension our hypothesis is consistent with clinical datasets, especially [24]. Their dataset includes *MYCN* mRNA and MYCN protein levels, presented as a Boolean variable (high or low) and a discrete variable with four possible values respectively. In order to compare our simulation results with their dataset, we set up comparable quantities within our classification system.

Working with the results of the first grid search, we noticed that the WT equilibria (*g_MYCN_* = 1) are always associated with less *MYCN* mRNA than their corresponding MA equilibria (*g_MYCN_* = 2). The mean value of the maximum and minimum *MYCN* mRNA levels in the combined set of WT and MA equilibria was used to define a critical threshold. An equilibrium above it was classified as having a high *MYCN* mRNA level and *vice versa*, in alignment with [24].

Working with the same set of simulation results, we merged the two sets of equilibrium MYCN protein levels (*g_MYCN_* = 1 and *g_MYCN_* = 2) and set up three thresholds to divide them into four intervals. They were selected to keep the 1259715 WT equilibria (*g_MYCN_* = 1) within the first three intervals and the 1259715 MA (*g_MYCN_* = 2) ones within the last three intervals, again in alignment with [24].

This binary classification system predicts a good outcome if p53/MYCN≥3.5 and a poor outcome otherwise. It also describes a tumour along three dimensions: its *MYCN* amplification status, *MYCN* mRNA level, and MYCN protein level. Along these dimensions, there are two (WT or MA), two (high or low), and four possible discrete values respectively: 16 categories in total. Our classification system reduces three dimensions to one, but the original three dimensions enable comparison between different datasets. We created this classification system partly by aligning the results of the first grid search with [24]. Although the thresholds (except the one for p53/MYCN) were manually imposed to achieve alignment, the mere existence of suitable thresholds is a plus point for our hypothesis.

### 2.4 Data mining and visualisation

To find evidence that our hypothesis is potentially correct in more general scenarios, we interpreted the simulation results of the three experiments within the classification system established on the basis of our hypothesis.

Three-dimensional scatter plots were created to visualise the results in terms of the *MYCN* mRNA levels (pM), MYCN protein levels (pM), and p53/MYCN metric levels (dimensionless) at the simulated equilibria.

The equilibria found in the second experiment were divided into four subsets along two dimensions. In one dimension, they were classified as WT or MA according to *g_MYCN_*. In the other dimension, they were classified as clinically good or poor according to the p53/MYCN metric. In each subset, we identified frequent itemsets of the tested parametric values and association rules between them with the Apriori algorithm [97]. The algorithm was implemented with the arules R package [98]. The technical details of this analysis are provided in subsection S2.8 in the supplementary file.

The equilibria found in the third experiment were also divided into the same four subsets. Although we could have applied the Apriori algorithm again after dividing each parameter’s range into discrete intervals, we decided to visualise this dataset directly by generating augmented scatterplot matrices. They illustrate how the equilibria in each subset are continuously distributed in the parametric space. How we used the ggpairs function within the GGally R package for this purpose is explained in subsection S2.8 in the supplementary file.

## 3 Results

This section begins by confirming the model’s stable long-term behaviour, presenting the results of a local sensitivity analysis (first experiment), and establishing the NB classification system. Then, the results of a global sensitivity analysis (second experiment) are presented within this classification system. After that, a more technical subsection delves deeper into the results of the second experiment by presenting the frequent itemsets and association rules extracted by the Apriori algorithm. The last subsection presents the results we produced in the third experiment.

In all figures, *MYCN_m_*, *MYCN*, and p53/MYCN indicate the *MYCN* mRNA level (pM), MYCN protein level (pM), and the p53/MYCN metric (dimensionless), respectively. A blue data point represents a WT equilibrium, whilst a red data point represents a MA equilibrium.

### 3.1 Stability analysis, local sensitivity analysis, and binary NB classification system

In the first experiment, we tested over 2.5 million parametric combinations, including two *MYCN* amplification statuses (*g_MYCN_* = 1 and *g_MYCN_* = 2). In each case, the state vector equilibrated to the same eight-dimensional fixed point from 10 different initial conditions. Furthermore, all of them were found to be stable. The results suggest that periodic, quasi-periodic, and chaotic dynamics are not biologically relevant in this context even if they can in theory be reached from certain basins of attraction when given certain parametric combinations. Therefore, our model complies with the requirement that oncogenic mutations abolish p53 oscillations [32].

A subset of the same dataset is presented in figure 5. Recall that there are three groups of parameters: the six stressors, *h*_5_, and the transcription and translation rates. Each panel presents the fixed points achieved by varying one group while keeping the other two at the default values, with or without *MYCN* amplification. The left panels indicate that the stressors principally affect the p53 protein level. The middle panels contain results that, relative to the results in the left and right panels, suggest that the model is not sensitive to *h*_5_. The fixed points in the right panels demonstrate large variations relative to the variations in the other four panels.

**Fig. 5:**
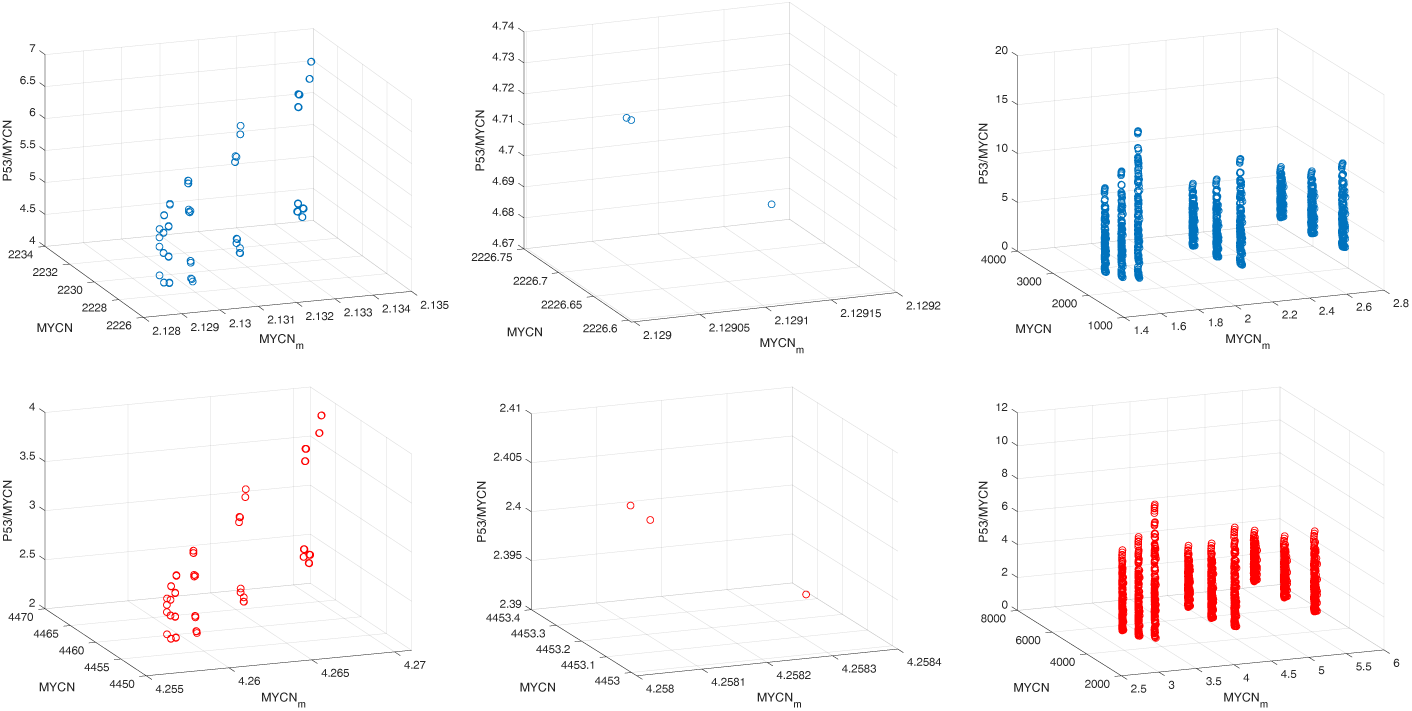
3D visualisation of the fixed points reached in the local sensitivity analysis. They are a subset of the results of the first experiment. The left panels present the fixed points obtained by switching on/off the stressors. The middle panels present the fixed points obtained by varying *h*_5_ with *s*_1_ = 0.5, the other stressors switched off, and the rates set to the default values. Recall that *s*_1_ was not set to zero because it is relevant to *h*_5_. The right panels present the fixed points obtained by varying the transcription and translation rates. In total, 64 + 3 + 6561 = 6628 fixed points are presented for each of *g_MYCN_* = 1 and *g_MYCN_* = 2. A blue data point represents a WT equilibrium, whilst a red data point represents a MA equilibrium. Both concentrations are in pM, but the p53/MYCN metric is dimensionless.

Returning to the full dataset, we applied our hypothesis by setting thresholds in three dimensions. As a reminder, our hypothesis is that the p53 protein level in a NB tumour, relative to the MYCN protein level therein, decides whether it is likely to progress or regress. This metric is called p53/MYCN and a NB with low p53/MYCN would have a poor clinical outcome. A corollary of the hypothesis is that p53/MYCN should follow a concave function of *MYCN*’s mRNA level in a mixed collection of WT and MA NBs.

First, we focused on the two simulations with the default parameters and inactive stressors, one with *g_MYCN_* = 1 (WT NB) and one with *g_MYCN_* = 2 (MA NB). Using the p53 and MYCN protein levels at the two simulated equilibria, we calculated the p53/MYCN metric’s values there: approximately 4.526 for WT NB and approximately 2.483 for MA NB. Due to the fact that *MYCN* amplification is typically associated with poor outcomes, we calculated their arithmetic mean (approximately 3.5) and made this the critical threshold separating good outcomes (p53/MYCN≥ 3.5) from poor outcomes (p53/MYCN*<*3.5).

Second, a critical *MYCN* mRNA level set anywhere on [2.84, 2.98] would classify every MA fixed point in figure 6 as having a high *MYCN* mRNA level and every WT fixed point as having a low level, in agreement with [24]. The midpoint was chosen.

**Fig. 6:**
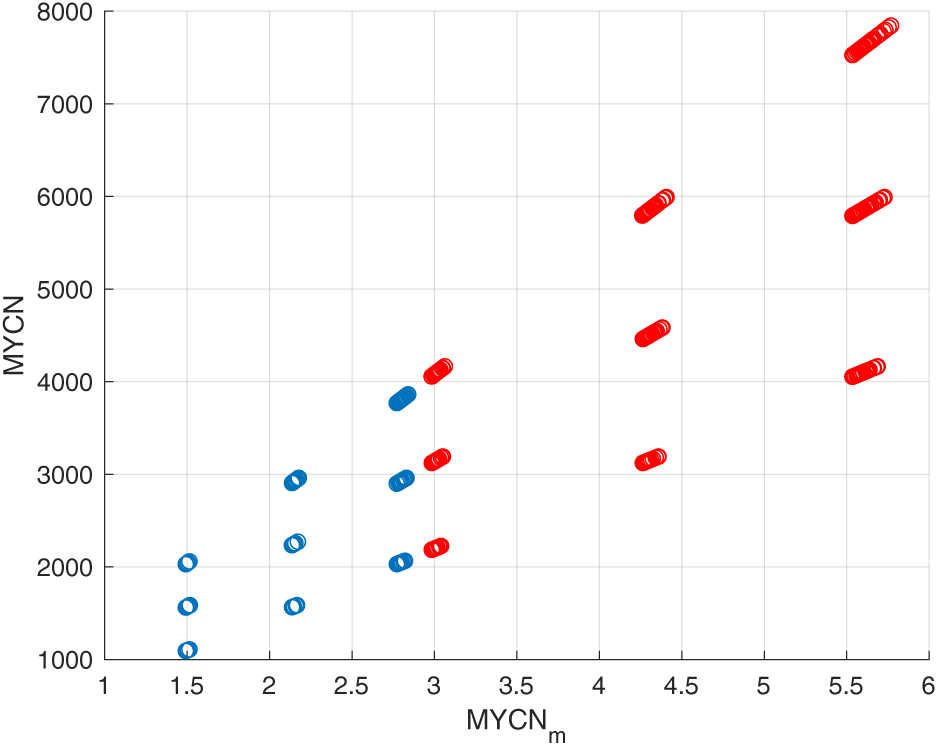
2D visualisation of the fixed points found in the first experiment. The x-axis reports *MYCN* mRNA levels (pM). The y-axis reports MYCN protein levels (pM). In this experiment, the stressors, *h*_5_, and the transcription and translation rates were varied together to generate 64 · 3 · 6561 + 3 = 1259715 parametric combinations. Each combination was used to perform two simulations, once with *g_MYCN_* = 1 and once with *g_MYCN_* = 2. A blue data point represents a WT (*g_MYCN_* = 1) equilibrium, whilst a red data point represents a MA (*g_MYCN_* = 2) equilibrium.

Third, the level of MYCN protein in [24] is a discrete variable with four possible values (zero, one, two, and three). Their WT levels are all smaller than or equal to level two, while their MA levels are all above level one. Like the level of *MYCN* mRNA, there are many solutions to this alignment problem, but we decided to respect the additional constraint of making the four intervals as evenly sized as possible. The following thresholds keep the simulated WT MYCN protein levels smaller than or equal to level two and the simulated MA levels greater than or equal to level one, with 88.89% of them being greater than or equal to level two.

- Level zero: from zero to 1500
- Level one: from 1500 to 2500.
- Level two: from 2500 to 4000.
- Level three: above 4000.

### 3.2 Correct predictions based on our hypothesis

Due to the results in the middle panel of figure 5, we decided to set *h*_5_ = 1 throughout the second experiment, but vary *s_i_* and the rates together. As a reminder, the range of p53/MYCN values in the middle panels is narrow relative to the ranges in the left and right panels in figure 5. Since six values of *g_MYCN_* (one, two, three, four, six, and eight) were used to mimic *MYCN* amplification, we ran 2519424 (64 · 6561 · 6) simulations.

Figure 7 presents the resulting fixed points within our classification system. The p53/MYCN metric predicts their clinical/treatment outcomes. Those above the critical threshold 3.5 are expected to have good clinical/treatment outcomes, whilst those below it are expected to have poor clinical/treatment outcomes.

**Fig. 7:**
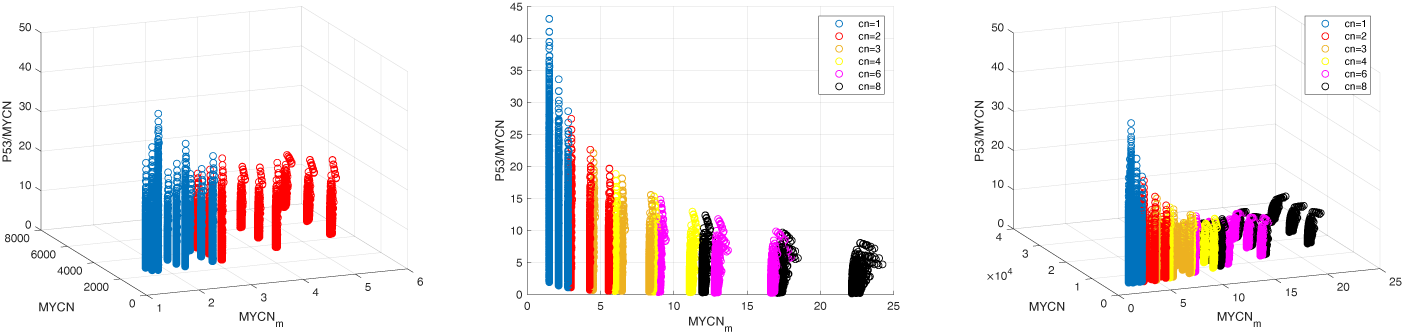
3D visualisation of the fixed points found in the second experiment, presented within our NB classification system. The stressors and rate parameters of the default configuration were varied together to produce 64 · 6561 = 419904 parametric combinations. Each combination was used to perform six simulations, with *g_MYCN_* set to one, two, three, four, six, and eight respectively. In the left panel, only the WT (*g_MYCN_* = 1, blue) and MA (*g_MYCN_* = 2, red) fixed points are presented. In the middle and right panels, the same fixed points are presented alongside the fixed points with even more copies of *MYCN* (three, four, six, and eight). Both concentrations are in pM, but the p53/MYCN metric is dimensionless.

Based on the p53/MYCN metric, 75.66 % of the WT fixed points (*g_MYCN_* = 1) belong to the category with good clinical/treatment outcomes. The remaining 24.34 % belong to the category with poor outcomes. The simulation results, when interpreted within our classification system and through the lens of our hypothesis, are in remarkably good agreement with [24]. Out of the 23 WT patients they studied, 18 achieved good outcomes (78.26 %) and five achieved poor outcomes (21.74 %).

Within our classification system, 32.03 % of the MA fixed points with *g_MYCN_* = 2 belong to the category with good outcomes. The remaining 67.97 % belong to the category with poor outcomes. Our results are aligned with the general clinical observation that an elevated *MYCN* copy number correlates with a progressively heightened prevalence of adverse clinical and biological characteristics [99]. These simulation results are also not too far from the specific measurements taken by [23, 24]. In Figure 2A in [23], although almost every *Trk* mRNA level reported for a MA specimen is close to zero (poor prognosis), three of the 15 reported levels are significantly higher, so 20% of the reported MA specimens achieved good outcomes. Figure 3C in [24] indicates that the six MA patients they studied all achieved poor outcomes.

Figure 7 also shows the effects of increasing *g_MYCN_* beyond two on p53/MYCN. The poor outcome percentage continues to increase with the copy number. For instance, with *g_MYCN_* = 3, 83.64% of the tested parametric combinations led to fixed points classified as poor outcomes. This percentage increases to 93.32%, 97,61%, and 98.78% with copy numbers 4, 6, and 8, respectively.

Along the other two dimensions (*MYCN* mRNA and MYCN protein levels), the simulation results and [24] are aligned within our classification system too. The simulated WT fixed points all have low *MYCN* mRNA levels, similar to [24]. The simulated MA fixed points with poor outcomes (67.97 %) all have high *MYCN* mRNA levels, just like Figure 3 in [24]. Table 3 compares the same pair of datasets along the third dimension, the protein level of MYCN. The trends are similar.

**Table 3:** Comparison of the simulation results (Sim) of the second experiment and [24] (Exp) along one dimension of our classification system: the level of MYCN protein. The simulation results are all fixed points, whilst the experimental results describe tumours. In this table, both datasets are divided into three categories based on the *MYCN* amplification status and outcome of a case. WT denotes a WT category (*g_MYCN_* = 1) and MA denotes a MA category (*g_MYCN_* = 2). The outcome is/was either good or poor. For example, 14.57 % of the WT fixed points with good clinical/treatment outcomes have MYCN protein levels that belong to level 0.

| Category | Level 0 | Level 1 | Level 2 | Level 3 |
| --- | --- | --- | --- | --- |
| WT-Good-Sim | 14.57 % | 63.57 % | 21.75 % | 0 |
| WT-Good-Exp | 22.22 % | 72.22 % | 5.56 % | 0 |
| WT-Poor-Sim | 0.02 % | 30.63 % | 69.35 % | 0 |
| WT-Poor-Exp | 0 | 0 | 100 % | 0 |
| MA-Poor-Sim | 0 | 3.52 % | 16.84 % | 79.63 % |
| MA-Poor-Exp | 0 | 0 | 33.33 % | 66.66 % |

Since we set the thresholds for the *MYCN* mRNA and MYCN protein levels by comparing the results of our first experiment with [24], the fact that our model agrees with [24] is not a genuine emergent phenomenon. However, our classification system’s ability to prognose NB cases based on the p53/MYCN metric is an emergent property. The critical threshold of p53/MYCN was defined independently of [24] and at no point were the patients’ clinical outcomes in [24] used in our procedures.

### 3.3 *MYCN* amplification produces diminishing returns in terms of p53/MYCN

In the second experiment, we increased *g_MYCN_* beyond two. The results are presented in the middle and right panels of figure 7. There is a clear trend that a higher copy number of *MYCN* is associated with lower and less variable values of p53/MYCN (consistently poor outcomes). The mean and standard deviation (SD) of this metric both decrease as *g_MYCN_* increases in figure 7. Numerically, the mean±SD (pM) is 6.04±3.63 with *g_MYCN_* = 1, 3.19±1.95 with *g_MYCN_* = 2, 2.24±1.40 with *g_MYCN_* = 3, 1.76±1.13 with *g_MYCN_* = 4, 1.27±0.85 with *g_MYCN_* = 6, and 1.02±0.71 with *g_MYCN_* = 8.

This decreasing trend is consistent with the clinically observed trend that in general, as the copy number of *MYCN* goes up in a tumour, its clinical and biological characteristics get progressively more adverse [99]. More specifically, the WT tumours studied by [23] displayed a much broader range of *TrkA* expression levels than their MA tumours, most of whose *Trk* mRNA levels were found to be close to zero. As the level of *TrkA* mRNA is a favourable prognostic indicator, just like the p53/MYCN metric (but mechanistically less relevant), this agreement exemplifies our classification system’s predictive ability. By extension, it is favourable for our hypothesis.

It is noteworthy that the p53/MYCN metric’s relationship with *g_MYCN_* is non-linear in the middle and right panels of figure 7, demonstrating diminishing returns. As the copy number increases from one to two, three, four, six and eight, the marginal loss in p53/MYCN decreases progressively. This agrees with a corollary of our hypothesis: the mRNA level of *MYCN* should have a non-linear relationship with p53 and MYCN levels.

Compared to the p53/MYCN metric, the levels of *MYCN* mRNA and MYCN protein vary with *g_MYCN_* in the opposite direction. The four means and standard deviations all increase with *g_MYCN_*. Consistently, [23] observed that their MA tumours almost uniformly expressed more *MYCN* mRNA than their WT tumours.

However, in our simulations, the general trend (the mean) in the relationship between the p53/MYCN metric and the mRNA level of *MYCN* is a monotonically decreasing function. Also, a low copy number of *MYCN* is associated with a broad range of p53/MYCN values. Our results disagree with the concave function suggested by figure 2A in [23].

### 3.4 Association rules extracted from the results of the second experiment

We attempted to gain mechanistic insights into the results obtained in the second experiment. To do so, we divided the fixed points into four subsets corresponding to four genotype-outcome combinations: WT (*g_MYCN_* = 1) or MA (*g_MYCN_* = 2) and good (p53/MYCN above 3.5) or poor (p53/MYCN below 3.5) clinical/treatment outcomes. We applied the Apriori algorithm to the parametric combinations that gave us the fixed points in each subset. Altogether, we extracted 80 association rules, 20 rules for each genotype-outcome combination. For the sake of clarity and conciseness, table 4 reports four meta association rules, which summarise the association rules. The four sets of 20 association rules are presented in tables S1, S2, S3, and S4 in the supplementary file. It is important to bear in mind that the association rules are robust but not very common. For example, in table S1, the top association rule is present in 1.1 % of the parametric combinations only (level of support). However, it is robust because of its high levels of confidence and lift (1 and 2.39), which clearly indicate that the antecedent (LHS) implies the consequent (RHS).

**Table 4:** Four meta association rules summarising the association rules extracted from four subsets of parametric combinations (second experiment). They describe wild-type and *MYCN* -amplified fixed points with good and poor outcomes. Each row presents the meta association rule summarising the dominant trait in 20 association rules. ‘/’ denotes OR. For example, H/High*MYCN* denotes H*MYCN* or High*MYCN*. Each rule follows the form: LHS ⇒ RHS, meaning that the left-hand side implies the right-hand side. L*X*, M*X*, H*X* denote the transcription rate of *X* is at 70%, 100%, 130% of the default value, respectively. Low*X*, Mid*X*, High*X* indicate the same for its translation rate.

| Genotype | Outcome | LHS | RHS |
| --- | --- | --- | --- |
| WT | Good | H/High*MYCN* and L/Low*p53* | L/Low*MYCN* |
| WT | Poor | H/High*p53* and s3on | L/Low*p53* |
| MA | Good | H/High*MYCN* and M/Mid*p53* | H/High*p53* |
| MA | Poor | H/High*p53* and L/Low*MYCN* | L/Low*p53* |

In the WT cases with good outcomes (table S1), the meta association rule is that a low *MYCN* transcription or translation rate is required when the tumour already exhibits a high *MYCN* translation or transcription rate and a low *p53* rate. Two rules indicate that when *MYCN*’s transcription and translation rates are both high and one of *p53*’s biosynthetic rate is at the default value, the other biosynthetic rate should be high (level of confidence is 0.97).

In the WT cases with poor outcomes (table S2), the meta association rule is that a low *p53* transcription or translation rate is required when the other *p53* biosynthetic rate is already high and the stressor *s*_3_ is active. Recall that the latter (*s*_3_) prevents MDM2 and p53 from forming complex C, where p53 degradation is enhanced. ATM is one protein that can perform this function in response to DNA damage (stress).

The MA fixed points with good outcomes (table S3) have very different association rules in their parametric combinations. 18 of the top 20 association rules pair a high *MYCN* biosynthetic rate and a default *p53* biosynthetic rate with another high *p53* biosynthetic rate.

The MA fixed points with poor outcomes (table S4) have less surprising association rules in their parametric combinations. 18 of the top 20 association rules pair a high *p53* transcription or translation rate and a low *MYCN* biosynthetic rate with a low *p53* translation or transcription rate. The antecedent also has one of the following: the other *MYCN* biosynthetic rate at the default value, an active *s*_3_, an active *s*_6_, an inactive *s*_4_, and a high *ARF* translation rate.

### 3.5 Random exploration of the parametric space

To test the robustness of the conclusions drawn from the first two experiments, the parametric space was explored randomly in the third experiment. 10000 random parametric combinations were created and each was used to run six simulations with different values of *g_MYCN_*. The resulting fixed points are shown in figure 8.

**Fig. 8:**
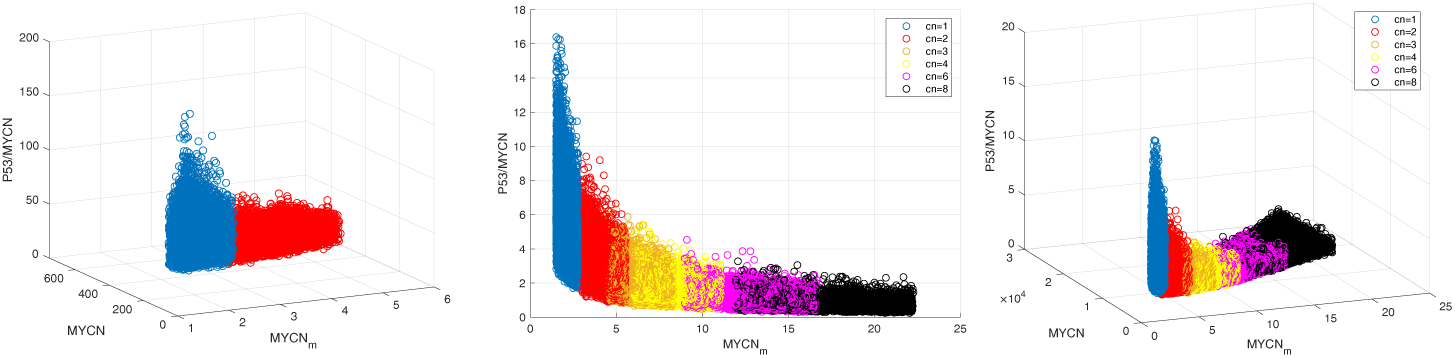
3D visualisation of the fixed points reached in the simulations in the third experiment. The stressors and rate parameters of the default configuration were varied together and randomly, whilst keeping *h*_5_ = 1, leading to 10000 parametric combinations. Each of these combinations was generated by sampling each stressor and rate randomly and uniformly between the minimum and maximum values tested in the second experiment. Each combination was used to perform six simulations, with *g_MYCN_* set to one, two, three, four, six, and eight respectively. In the left panel, only the WT (*g_MYCN_* = 1, blue) and MA (*g_MYCN_* = 2, red) fixed points are presented. In the middle and right panels, the same fixed points are presented alongside the fixed points with even more copies of *MYCN* (three, four, six, and eight). Both concentrations are in pM, but the p53/MYCN metric is dimensionless.

In figure 8, based on the p53/MYCN metric, 83.17% of the 10000 WT fixed points (*g_MYCN_* = 1) have good outcomes and the remaining 16.83% have poor outcomes. On the other hand, only 24.17% of the 10000 MA fixed points (*g_MYCN_* = 2) have good outcomes and the remaining 75.83% have poor outcomes. [24] studied 23 WT patients, 18 of whom had good outcomes (78.26%) and five had poor outcomes (21.74%). They only studied six MA patients and they all had poor outcomes (100%). Echoing the comments made in subsection 3.2, our results agree with the general clinical observation that an elevated *MYCN* copy number correlates with a poor clinical outcome [99].

The WT fixed points all have low levels of *MYCN* mRNA, just like [24]. The MA fixed points all have high levels of *MYCN* mRNA, just like [24]. As can be seen in table 5, the simulated MYCN protein levels mostly agree with [24] too.

**Table 5:** Comparison of the simulation results (Sim) of the third experiment and [24] (Exp) along one dimension of our classification system: the level of MYCN protein. The simulation results are all fixed points, whilst the experimental results describe tumours. In this table, both datasets are divided into three categories based on the *MYCN* amplification status and outcome of a case. WT denotes a WT category (*g_MYCN_* = 1) and MA denotes a MA category (*g_MYCN_* = 2). The outcome is/was either good or bad. For example, 10.67 % of the WT fixed points with good clinical/treatment outcomes have MYCN protein levels that belong to level 0.

| Category | Level 0 | Level 1 | Level 2 | Level 3 |
| --- | --- | --- | --- | --- |
| WT-Good-Sim | 10.67 % | 67.52 % | 21.81 % | 0 |
| WT-Good-Exp | 22.22 % | 72.22 % | 5.56 % | 0 |
| WT-Poor-Sim | 0 | 33.1 % | 66.9 % | 0 |
| WT-Poor-Exp | 0 | 0 | 100 % | 0 |
| MA-Poor-Sim | 0 | 0.3 % | 27.48 % | 72.21 % |
| MA-Poor-Exp | 0 | 0 | 33.33 % | 66.66 % |

The results of increasing *g_MYCN_* beyond two are presented in the middle and right panels of figure 8. There is a clear trend that a higher copy number of *MYCN* is associated with lower and less variable values of p53/MYCN (consistently bad outcomes). The mean and SD of this metric both decrease as *g_MYCN_* increases in figure 8: 5.54±2.11 (mean±SD) with *g_MYCN_* = 1, 2.88±1.11 with *g_MYCN_* = 2, 1.98±0.75 with *g_MYCN_* = 3, 1.57±0.58 with *g_MYCN_* = 4, 1.14±0.43 with *g_MYCN_* = 6, and 0.90±0.34 with *g_MYCN_* = 8. Reflecting the decreasing trend of the p53/MYCN metric’s mean, 95.70% of the fixed points with *g_MYCN_* = 3 have poor outcomes; *g_MYCN_* = 4, 99.28%; *g_MYCN_* = 6, 99.94%; and *g_MYCN_* = 8, 100%. Compared to the p53/MYCN metric, the levels of *MYCN* mRNA and MYCN protein vary with *g_MYCN_* in the opposite direction. The two means and two standard deviations all increase with *g_MYCN_*.

These results mirror those presented in figure 7, so the comments made in subsection 3.3 are true for the results presented in figure 8 too. In agreement with [23], the WT fixed points display a much broader range of p53/MYCN than the MA fixed points. p53/MYCN displays a non-linear relationship with *g_MYCN_*. Aligned with [23], the levels of *MYCN* mRNA and MYCN protein become higher when *g_MYCN_* increases in figure 8. Again in conflict with figure 2A in [23], p53/MYCN is a monotonically decreasing function of the mRNA level of *MYCN* and p53/MYCN is highly variable when *g_MYCN_* is small.

### 3.6 Visualising parametric distributions in the four scenarios of the third experiment

Just as we did before learning association rules from the results of the second experiment, we divided the fixed points found in the third experiment into four subsets (genotype-outcome combinations): WT (*g_MYCN_* = 1) or MA (*g_MYCN_* = 2) and good (p53/MYCN metric above 3.5) or poor (p53/MYCN metric below 3.5) clinical/treatment outcomes. Figure 9 is a set of four ggplot2 generalised pairs plots. They visualise the four subsets of parametric combinations. Each plot was created using the ggpairs function, publicly available in the GGally R package.

**Fig. 9:**
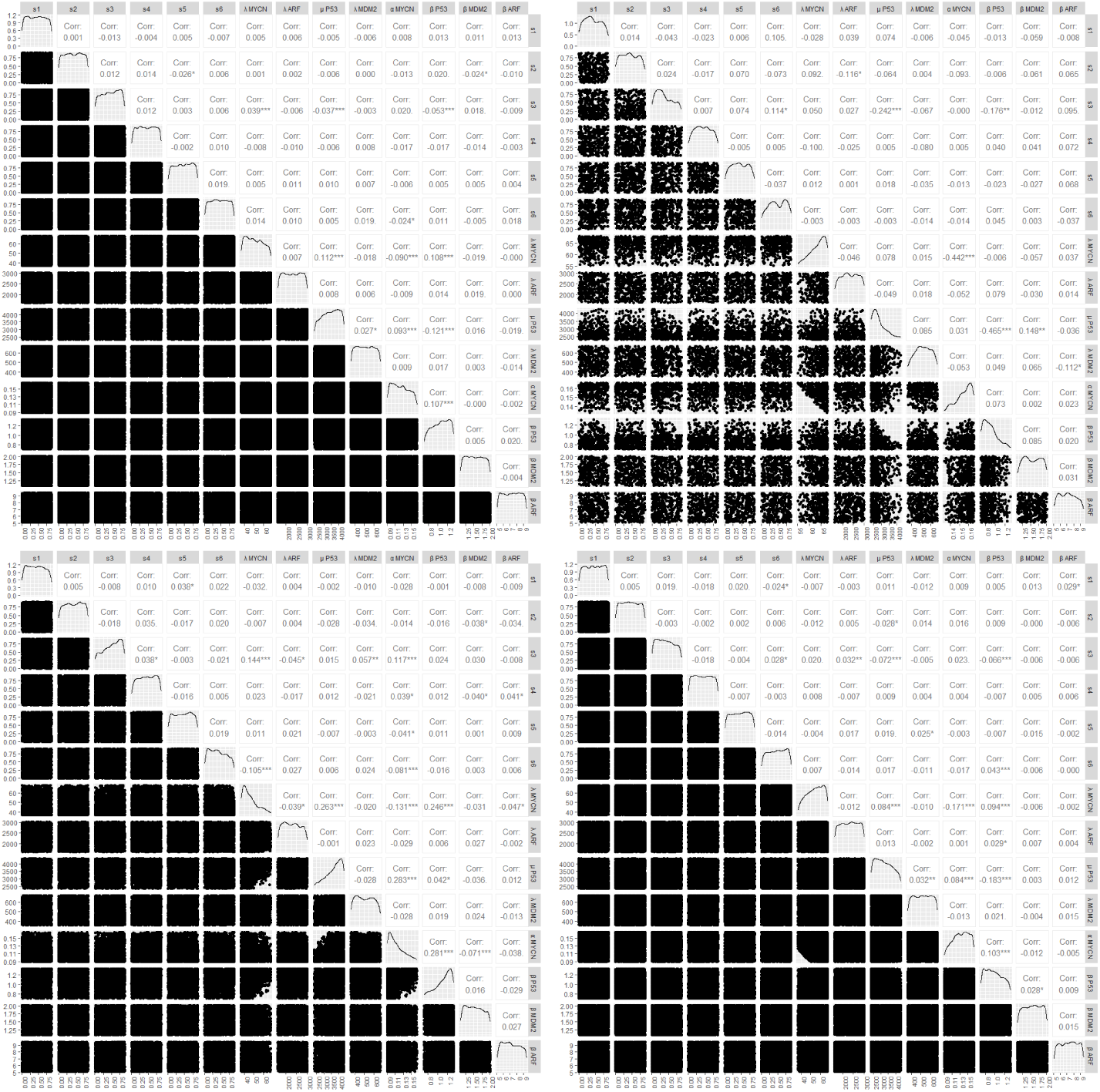
Four ggplot2 generalised pairs plots comparing pairs of 14 parameters (six stressors and eight rates) in four subsets of the parametric combinations tested in the third experiment. The top two plots present the parametric combinations leading to WT fixed points. The bottom two plots present the parametric combinations leading to MA fixed points. The left two plots present the fixed points with good outcomes. The right two plots present the fixed points with poor outcomes. In each plot below the leading diagonal are 91 scatterplots comparing the unique pairs formed by the 14 parameters. On the diagonal are the probability density functions of the 14 parameters in the subset. Above the diagonal are the Pearson correlation coefficients of the 91 pairs of parameters. Each correlation’s statistical significance is shown too. In a matrix element, *** means the reported p-value is smaller than 0.001; ** means that it is smaller than 0.01; * means that it is smaller than 0.05; and a full stop (.) means it is smaller than 0.10.

The top-left panel in figure 9 reveals interesting features shared by the parametric combinations giving us WT fixed points with good outcomes. The association rules in table S1 are robust but rare. They indicate the requirements (RHS) for such a fixed point given a set of conditions (LHS), but the conditions are rare. By contrast, this plot pertains to each parameter’s distribution in this sample, including its average value. The two distributions of *MYCN* rates are skewed towards their minima, whilst both distributions of *p53* rates are skewed towards their maxima. The six stressors have relatively uniform distributions in this subset, implying that having specific combinations of stressors is unnecessary for a good outcome.

The top-right panel pertains to the WT fixed points with poor outcomes. The distributions of the two *MYCN* rates are skewed towards their maxima, whilst the distributions of the two *p53* rates are skewed towards their minima. Furthermore, the plot indicates that it is impossible for both of *MYCN*’s rates to be low or both of *p53*’s rates to be high in a parametric combination. Interestingly, the distribution of *s*_3_ is skewed towards its minimum.

The bottom-left panel pertains to the MA fixed points with good outcomes. The distributions of both *MYCN* rates are skewed towards their minima, whilst the distributions of both *p53* rates are skewed towards their maxima. Furthermore, it is impossible for both *MYCN* rates to be very high or for a low *p53* rate to coexist with a high *MYCN* rate in a parametric combination. Compared to the top-left panel (WT, good outcomes), where the six stressors are almost uniformly distributed, *s*_3_ and *s*_6_ are skewed towards the maximum and minimum respectively.

The bottom-right panel presents the parametric combinations giving us MA fixed points with poor outcomes. The distributions of both *MYCN* rates are skewed towards their maxima, but the distributions of both *p53* rates are skewed towards their minima. It is impossible for both *MYCN* rates to be very low in a parametric combination.

## 4 Discussion

The *MYCN* enigma is the general observation that a NB patient’s clinical outcome does not depend monotonically on their tumour’s *MYCN* expression level or *MYCN* amplification status. This paper presents a hypothesis inspired by MYCN’s ability to activate *p53* despite the consensus that *MYCN* is an oncogene [21] and *p53* is a tumour suppressor gene [25]. The key concept in this hypothesis is that the p53 protein level in a NB tumour, relative to the MYCN protein level therein, decides whether it progresses or regresses.

We built a mathematical model comprising eight non-linear ODEs to model a molecular network connecting MYCN to the ARF/MDM2/p53 axis. The model itself is an addition to the literature on NB for the following reasons. The negative feedback loop formed by MDM2 and p53 has been modelled with ODEs in many studies, mostly with the aim of understanding how genetic damage triggers oscillations in p53’s level [32–38]. Some models even couple p53 dynamics with ARF and other species acting upstream (such as ATM) and downstream (such as WIP1) of p53 [39–43]. However, to the best of our knowledge, the presented mathematical model is the only one that couples MYCN to the ARF/MDM2/p53 axis.

After parameterising the model, we tested the model to confirm that its long-term behaviour is stable for over 1.25 million parametric combinations with and without *MYCN* amplification. Only stable fixed points were found, so our model complies with [32]. Their theory states that oncogenic mutations abolish p53 oscillations. Then, we explored the parametric space systematically on a grid. We varied the pattern of active stressors (up to six), the Hill coefficient *h*_5_, and the eight transcription and translation rates. We varied the three groups independently and together at different levels of *MYCN* amplification, including the wild type without this mutation. Finally, we explored the parametric space randomly.

### 4.1 To what extent is the hypothesis correct?

We applied our hypothesis to the simulation results of the first experiment to establish thresholds along three dimensions: *MYCN* mRNA’s level, MYCN protein’s level, and the p53/MYCN metric. This turned our hypothesis into a concrete binary classification system. It predicts whether a patient is likely to have a good or poor outcome based on the p53/MYCN metric level in their tumour. It also describes a tumour along three dimensions: its *MYCN* amplification status (Boolean variable), *MYCN* mRNA level (Boolean variable), and MYCN protein level (discrete variable with four possible values). These extra dimensions enable comparison between simulated fixed points and real tumours.

Figure 5 presents the fixed points found in the first experiment. Compared to the six stressors and eight transcription/translation rates, the Hill coefficient *h*_5_ is not a sensitive parameter. It controls the importance of cooperativity in *s*_1_’s positive influence on *ARF* transcription. *s*_1_ models oncogenic stresses that activate *ARF* transcription through E2F-1 [60–62] and Ras [63, 64], as well as oxidative and heat shock stresses [65]. Unlike the other species, E2F-1 is a transcription factor, so it physically interacts with DNA and *h*_5_ is most relevant to this aspect of *s*_1_. Therefore, our simulation results suggest that cooperativity plays a small role in how E2F-1 influences p53/MYCN.

Interpreted within this classification system, the results of the second experiment (figure 7 and table 3) are consistent with most of the clinical and experimental observations listed in section S1 in the supplementary file.

- Figure 7 clearly indicates a negative relationship between the copy number of *MYCN* and the p53/MYCN metric, a favourable biomarker in our classification system, suggesting a tumour’s clinical and biological characteristics get progressively more adverse with its degree of *MYCN* amplification. This is consistent with [99]. The non-linear relationship in figure 7 is a corollary that follows from our hypothesis too. When the *MYCN* copy number is higher, the marginal change to p53/MYCN associated with an additional copy of *MYCN* is lower.
- The p53/MYCN metric correctly predicts positive outcomes for around three quarters of the wild-type fixed points, just like the 23 WT patients [24] studied.
- The p53/MYCN metric indicates that most of the MA fixed points have poor outcomes. Three of the 15 MA patients studied by [23] and all six of the MA patients studied by [24] achieved poor outcomes. The discrepancies could be explained by two reasons. First, the two datasets [23, 24] suffer from small sample sizes. Second, our results pertain to a *MYCN* copy number of two only and their MA patients might have acquired more copies of *MYCN*. Indeed, in figure 7, the poor outcome percentage continues to increase when the copy number increases beyond two.
- Within our classification system, the simulated WT and MA fixed points all have low and high *MYCN* mRNA levels respectively, just like their clinical counterparts [24]. This trend is consistent with the observations of [23] too. Almost uniformly higher *MYCN* mRNA levels were measured in their MA specimens than their WT specimens. Table 3 indicates that the simulated protein levels match [24] very closely.
- In figure 7, there is a clear trend that a higher copy number of *MYCN* is associated with not just lower but also less variable values of p53/MYCN (consistently bad outcomes).
- Interestingly, figure 7 does not resemble the graph of a concave function like Figure 2A in [23]. When the level of *MYCN* mRNA is very low, the p53/MYCN metric displays a broad range of values. To be aligned with [23], the WT fixed points with inadequate MYCN should all be under the critical threshold of 3.5. Conversely, the WT fixed points with enough MYCN should all be above 3.5. One possibility is that WT NB tumours have a mechanism (refer to subsection 4.3) triggered by a low level of MYCN to keep p53/MYCN under 3.5. It follows that the left-most blue fixed points above 3.5 in the middle panel of figure 7 are biologically irrelevant. Another explanation is that [23] investigated only seven WT cases (small sample size).

### 4.2 Expanding the hypothesis by going beyond the model

Briefly, the results discussed in the last subsection support the conclusion that upon *MYCN* amplification, it becomes more difficult for a NB tumour to have p53/MYCN over 3.5. The conclusion favours our hypothesis as an explanation for the clinical and experimental results constituting the *MYCN* enigma. A patient’s clinical outcome depends neither on *MYCN* amplification nor *MYCN* expression alone. The level of p53/MYCN is the key.

However, we remain ignorant of why it works and our hypothesis is clearly incomplete. It does not explain why the p53/MYCN metric should be kept under 3.5 in most WT tumours with inadequate MYCN and *vice versa*. This subsection aims to fill these gaps by examining the features shared by each group of parametric combinations that gave us a particular category of fixed points (genotype-outcome combination) in the second and third experiments. As a reminder, these features are the four sets of association rules in section S3 in the supplementary file and the parametric distributions in figure 9. We will do so by speculating about what a real NB cell does to create the necessary conditions (parametric values). These speculations will take us beyond the scope of the mathematical model.

In the WT cases with good outcomes (table S1), the meta association rule implies that competitive dynamics are involved. When *MYCN* transcription is elevated, it takes resources away from *MYCN* translation and *vice versa*. That *p53*’s transcription or translation rate is low in 18 of the top 20 association rules suggests that the biosynthetic processes producing MYCN and p53 compete for the same resources. It is true that two rules suggest that *MYCN*’s transcription and translation rates can both be high, but they are exceptions rather than the norm. For the parametric combinations giving us WT fixed points with good outcomes in the third experiment (top-left panel, figure 9), the two distributions of *MYCN* rates are skewed towards their minima, whilst both distributions of *p53* rates are skewed towards their maxima. These features are consistent with the need to produce more p53 than MYCN to keep p53/MYCN high (good outcome). One possibility is that a transcription factor upregulates biogenesis in general, but it preferentially boosts *p53*’s processes. The relatively uniform distributions of the six stressors are consistent with a mechanism where these regulatory processes are independent of the six stressors, mirroring the predominantly good clinical outcomes observed in WT NB cases. We further hypothesise that MYCN itself is the regulator. It follows from our prediction that in a real WT tumour, the biosynthetic rates in these parametric combinations should not be possible when there is not enough MYCN. In figures 7 and 8, some of the blue points with p53/MYCN above 3.5 and MYCN at low levels are probably biologically irrelevant.

In the WT cases with poor outcomes (table S2), the meta association rule implies that the processes of *p53* transcription and translation compete for the same resources. The distributions of the four *MYCN* and *p53* rates (top-right panel, figure 9) are all consistent with the need to produce MYCN more efficiently than p53 to keep p53/MYCN low (poor outcome). *s*_3_ is active and *p53* has a high rate in all 20 association rules, whilst the distribution of *s*_3_ in the whole sample is skewed towards its minimum. One possibility is that when MYCN is scarce, ATM signalling (or a functionally equivalent pathway) is triggered to restore biogenesis in general, but it preferentially boosts MYCN production. If ATM signalling gets too high, p53 production will increase too (association rules and [100]), but MYCN production will still be higher to maintain the WT-poor scenario. When there is enough MYCN, ATM signalling shuts down and MYCN becomes responsible for biogenesis again (back to the WT-good scenario). In addition to explaining why WT tumours have poor outcomes when they lack MYCN, our hypothesis also predicts that the biosynthetic rates giving us WT fixed points with poor outcomes should not be possible when there is enough MYCN (ATM signalling is off). In figures 7 and 8, some of the blue points with p53/MYCN below 3.5 and MYCN at relatively high levels may be biologically irrelevant.

The distributions of *MYCN*’s and *p53*’s rates in the subset of parametric combintions giving us MA fixed points with good outcomes (bottom-left panel, figure 9) are as expected. A MA tumour must produce p53 more efficiently than it produces MYCN to achieve a good outcome. However, the meta association rule extracted from this subset (table S3) suggests that MA patients require extremely high biosynthetic rates to achieve good outcomes. We speculate that even MA tumours with abundant MYCN rarely achieve the required rates. The distributions of *s*_3_ and *s*_6_ are skewed towards the maximum and minimum respectively, whilst the corresponding distributions in the WT cases with good outcomes (top-left panel, figure 9) are almost uniform. The implication is that upon *MYCN* amplification, only certain combinations of active stressors can result in a good outcome. Recall our prediction made in the previous paragraph that when there is enough MYCN (as in a MA tumour), ATM signalling (a part of *s*_3_) should become inactive. The distribution of *s*_3_ in the bottom-left panel of figure 9 contradicts this prediction. We speculate that MA tumours with abundant MYCN rarely achieve the combinations of active stressors required for good clinical outcomes. Specifically, they cannot keep ATM signalling on. It follows that the red points (and others that are not blue) with p53/MYCN above 3.5 in figures 7 and 8 are biologically rare even if they are possible.

The distributions in the bottom-right panel of figure 9 (MA, poor outcomes) are aligned with the intuition that a MA tumour must produce MYCN more efficiently than it produces p53 to achieve a poor outcome (p53/MYCN below 3.5). The almost uniform distributions of all six stressors are consistent with the consensus that MA tumours usually have poor outcomes, regardless of the combination of active stressors therein. We speculate that MYCN preferentially boosts *p53*’s rates in WT tumours and *MYCN*’s rates in MA tumours. Maybe, *MYCN* amplification occurs with a regulatory shift, shifting the preference from p53 production to MYCN production. *MYCN* amplification could be responsible for this shift, either directly by producing more MYCN or indirectly by triggering a stress response. Alternatively, a co-occurring mutation could be responsible. For example, *ALK* mutations frequently co-occur with *MYCN* amplification; activated ALK and MYCN collaborate in NB pathogenesis [101, 102].

### 4.3 Biological basis of the expanded hypothesis

In subsection 4.2, we went beyond the scope of our model to speculate on the mechanistic details of our hypothesis, including why our results differ from parts of [23] and [24]. The first component of the expanded hypothesis states that MYCN upregulates biogenesis in general in NB tumours. The second component predicts that ATM signalling is activated when MYCN is scarce to restore biogenesis in general and MYCN production in particular. When MYCN’s level is restored, ATM signalling becomes inactive again. The third component states that MA tumours require unrealistically high biosynthetic rates for good outcomes. These rates are hard to attain; so is the high level of ATM activity when MYCN is abundant. The final component is that there is a regulatory shift when *MYCN* amplification occurs, causing MYCN to shift from preferentially boosting p53 production to MYCN production. This expanded hypothesis is consistent with the NB literature, as discussed below.

First component. Although MYCN is known for its role as a transcription factor, it has numerous interaction partners. It interacts with different partners in different environments [103]. As reviewed by [103], MYCN interacts with DNA topoisomerase II proteins to resolve torsional stress in supercoiled DNA, prevents R-loop formation, resolves R-loops, interacts with the RNA exosome to degrade aberrant RNA species, and binds to RNA species directly to enhance stress resilience. These functions allow transcription to proceed efficiently. MA NB cells express high levels of PARP enzymes, which are activated at sites of DNA damage and upon activation, recruit repair factors [103]. A signalling pathway linking MYCN through PARP enzymes to DNA damage response (DDR) genes such as BRCA1 was experimentally identified by [104]. In the experiment [104], suppressing MYCN, PARP enzymes, and DDR genes led to reduced cancer cell survival. Finally, MYCN regulates protein synthesis [105] and folding [106], as well as cellular metabolism [103, 107]. These functions allow translation to proceed efficiently.

Second and third components. Considering MYCN counteracts so many stresses, including DNA damage (previous paragraph), it is not unreasonable to speculate that MYCN deficiency activates compensatory cellular stress responses to restore biogenesis. For example, the HSF1-mediated proteotoxic stress response is a transcriptional programme activated by misfolded proteins; it produces translation machinery components [108]. The integrated stress response activates an adaptive gene expression programme [109]. The ATM (a part of *s*_3_) and ATR DDR pathways are both promoted [110]. Interestingly, ATM signalling activates p53 [100], mirroring the 20 association rules in table S2, which state that *p53* has a high rate when *s*_3_ is active. This idea is even more appealing if NB cells are addicted to MYCN. Many cancers are addicted to oncogenes [111]. Addiction is likely in this particular case because despite DDR defects, NB tumours can tolerate high levels of stress thanks to MYCN’s actions [15]. For example, MYCN downregulates ATM by miR-421 [112] and repairs damaged DNA (previous paragraph). In fact, the first action is exactly what the second component predicts—when MYCN’s level is restored, ATM signalling becomes inactive again. Its existence provides a biological basis for the third component too—MA tumours have abundant MYCN and are unlikely to experience a high level of ATM activity.

Fourth component. In a MA tumour, which always has enough MYCN, MYCN upregulates biogenesis. Compared to a WT tumour, however, MYCN preferentially boosts MYCN production rather than p53 production in a MA tumour. This regulatory shift could be caused by *MYCN* amplification itself or by mutations that co-occur with *MYCN* amplification in neuroblastoma, perhaps in the *ALK* gene [101, 102]. This explanation is plausible because [113] experimentally demonstrated the ability of both WT ALK and gain-of-function ALK to initiate *MYCN* transcription.

### 4.4 Validity

In the first experiment, only stable fixed points were found as predicted by [32]. Using the model, we predicted two novel emergent phenomena, namely that E2F-1 does not transcribe *ARF* cooperatively and a non-linear relationship between the *MYCN* copy number of a patient and their outcome. Our model’s ability to reproduce and predict emergent phenomena makes it a contribution to the field in its own right.

The fact that our simulation results are consistent with most of the clinical and experimental observations listed in section S1 in the supplementary file is encouraging. Also reassuring is that searching the parametric space systematically (experiment two) and randomly (experiment three) gave us similar results. Our hypothesis that the p53/MYCN metric decides a NB tumour’s clinical/treatment outcome is valid and deserves further investigation.

Whilst our results differ from [23] and [24] in some respects, the expanded hypothesis proposed in subsection 4.2 reconciles the differences in theory. Furthermore, it is supported by the publications mentioned in subsection 4.3, so we believe that it deserves attention from experimental and clinical researchers.

Nevertheless, it must be acknowledged that the model’s structure and the chosen parametric values form an incomplete representation of how things work within neuroblastoma cells.

### 4.5 Future work

The four components of the expanded hypothesis, which we proposed in subsection 4.2 by adding mechanistic links not already in the mathematical model, are not testable even *in silico*. Therefore, it remains a tentative explanation of the *MYCN* enigma. For it to become a well-substantiated theory explaining how MYCN affects a heterogeneous population of NB cells by interacting, cooperating, and competing with p53 in different cellular contexts (combinations of mutations), we must build a more comprehensive mathematical model to guide an experimental study.

The improved model would replace the eight static biosynthetic rates with functions dependent on the protein level of MYCN and the mutations present in a NB cell. It would include a negative feedback loop between MYCN and *s*_3_ too. The static stressor would be replaced by a set of equations modelling a set of stress responses to MYCN deficiency. Based on subsection 4.3, the HSF1-mediated proteotoxic stress response [108], the integrated stress response [109], and various DDR pathways (including ATM and ATR signalling) [100, 110] are potential members of this set. Closing the loop, the new variables representing the selected stress responses would be connected to the biosynthetic rates.

More fundamentally, the p53/MYCN metric is a population-level metric. When the hypothesis refers to its level in a cell, it is actually the average cell in a NB tumour. The hypothesis assumes that p53 is pro-apoptosis in a population of identical NB cells. However, as we proposed recently [114], a MA tumour may contain two distinct populations of cancer cells, where p53 is pro-survival and pro-apoptosis respectively. The basis is that p53 regulates hundreds of genes in a context-dependent manner, so it has diverse and contradictory downstream effects, including DNA repair, cell cycle arrest, and apoptosis [115].

The updated mathematical model should be incorporated into each NB cell agent in the multicellular model presented in our recent publications [114, 116]. Currently, although MYCN and p53 are both multifunctional as required in the multicellular model and it naturally describes a heterogeneous population, the *MYCN* enigma is modelled simplistically as a conditional statement involving three Boolean variables. It states that if *MYCN* is amplified and MYCN is active in a NB cell agent, then p53 is inactive therein. The updated mathematical model, which would describe the molecular network more accurately, would represent the *MYCN* enigma better.

One could use the modified multicellular model to test the population-level p53/MYCN metric’s ability to predict clinical/treatment outcomes in different scenarios. p53 has different functions, which could be modelled as a probability distribution over a heterogeneous population of NB cells. This change would generalise the p53/MYCN metric and make it applicable at the cellular level of resolution.

## 5 Conclusion

The *MYCN* enigma—the observation that the outcome of neuroblastoma does not depend monotonically on *MYCN* expression or amplification—motivated our hypothesis that the amount of p53 protein relative to the amount of MYCN protein is a superior prognostic metric. Developing a novel mathematical model linking MYCN to the ARF/MDM2/p53 axis, we carried out numerical simulations to test our hypothesis. The results are aligned with numerous clinical and experimental observations.

After extracting association rules and parametric correlations from the tested parametric combinations, we expanded the hypothesis by adding mechanisms outside the scope of the mathematical model. According to the expanded hypothesis, MYCN is responsible for biogenesis in NB cells and *MYCN* amplification and/or co-occurring mutations trigger a regulatory shift to reduce p53/MYCN (worse outcome). The expanded hypothesis also predicts a negative feedback loop between MYCN deficiency and ATM signalling (or equivalent), which restores biogenesis and favours MYCN production in a WT tumour deficient in MYCN. Finally, it predicts that a MA tumour requires very high biosynthetic rates and ATM signalling (or equivalent) to achieve a good outcome (p53/MYCN above 3.5). These conditions are very hard to attain biologically.

The expanded hypothesis is supported by the literature, but biosynthesis and stress responses to MYCN deficiency are represented simplistically in the model. More mechanistic details about them must be added to the model. The multifunctional nature of p53 and tumour heterogeneity must be taken into account too. Such an improved model could enable more detailed, granular, and comprehensive simulations, which could in turn inform lab-based experiments and therapeutic design.

## 6 Data availability statement

This manuscript has no associated data. Everything is included in the main manuscript and supplementary information section.

## 7 Author contributions and acknowledgments

KYW conceived the project and secured funding. MP and KYW built the mathematical model with input from DW and RD. MI found the model parameters with input from KYW. KYW and MI proposed that p53/MYCN is a prognostic metric for neuroblastoma. MI designed and implemented the numerical simulations with input from KYW and FD. MI analysed the simulation results with input from KYW and FD. MP and MI created the visualisations with input from KYW and FD. KYW proposed the expanded hypothesis with input from MI, FD, and DW. KYW, MI, FD, and RD wrote the manuscript. KYW and DW supervised MP and RD. KYW and FD supervised MI. All six authors have read and approved the manuscript. KYW and RD are grateful for funding provided by the Insigneo Institute for *in Silico* Medicine. The authors want to thank Daniel Jordan for the preliminary results that inspired this project.

## Supporting Information

### S1 Clinical and experimental observations of relevance to the *MYCN* enigma

A satisfactory explanation of the *MYCN* enigma must be consistent with the following clinical and experimental observations.

- In general, as the copy number of *MYCN* goes up in a NB tumour, its clinical and biological characteristics get progressively more adverse to survival [99].
- Out of the 23 WT patients [24] studied, 18 achieved good outcomes (78.26 %) and five achieved poor outcomes (21.74 %). The six MA patients they studied all achieved poor outcomes. In another study [23], only three out of 15 MA patients achieved good outcomes (20 %) and the rest achieved poor outcomes (80 %). Together, the two datasets [23, 24] suggest that an average MA patient has a 14.29% chance of achieving a good outcome.
- [24] measured the levels of *MYCN* mRNA and MYCN protein in their patients’ tumours. The results are reported in Fig. 3C in their paper. Tumours with similar *MYCN* mRNA levels were found to have very different MYCN protein levels.
- [23] measured the levels of *MYCN* mRNA and *TrkA* mRNA in 76 WT primary NB specimens and 15 MA ones. The results are reported in Figure 2A in their paper. They measured almost uniformly higher *MYCN* mRNA levels in the MA specimens than the WT specimens. It should be noted that whilst *TrkA* expression is indeed a favourable biomarker for neuroblastoma patients, it does not follow that *TrkA* mechanistically suppresses neuroblastoma.
- Again in Figure 2A in [23], almost every *Trk* mRNA level reported for a MA specimen is close to zero (poor prognosis). The values reported for the WT specimens are much more diverse, showing a positive and linear relationship with the corresponding *MYCN* mRNA levels. Overall, the figure presents a concave function between the two sets of mRNA levels.
- NB cells are cancer cells. If the explanation involves *p53*, it must agree with a recent theory [32]. It states that oncogenic mutations abolish p53 oscillations.

### S2 Methodological details

#### S2.1 Master Equations

Our gene regulatory network comprises four genes. Each pair of master equations describes the concentration dynamics of a gene’s mRNA transcript and protein. Equation S1 models transcription in terms of the transcript’s production and degradation rates. Equation S2 describes translation in terms of the protein’s production and degradation rates. In other words, each equation describes phenomena occurring at two timescales, namely synthesis and degradation. The two equations are linked by the mRNA transcript’s concentration level, on which the translation rate depends.

The master equations for gene *i*, where *m_i_*(*t*) and *p_i_*(*t*) describe the concentration levels of its mRNA transcript and protein, are given below:

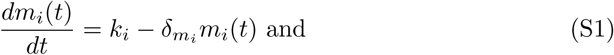

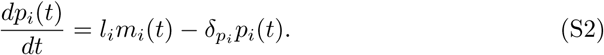

In equations S1 and S2, *k_i_* represents the transcription rate of gene *i* and *l_i_* represents its translation rate. In the same equations, *δ_mi_* and *δ_pi_* are the degradation rates of its mRNA transcript and protein respectively. After tailoring equations S1 and S2 to describe the four genes, we will use more readable variables than *m_i_*(*t*) and *p_i_*(*t*) to represent the eight concentrations (pM): [*MY CNm*] and [*MY CN*]; [*p*53*m*] and [*p*53]; [*MDM* 2*m*] and [*MDM* 2]; and [*ARFm*] and [*ARF*]. *MYCN* amplification can be simulated by increasing *k_MY CN_*.

The key property of this pair of equations is that they are both linear, but the model pertains to a non-linear problem, so it should be non-linear. The model is non-linear because it also describes non-linear binding events. First, stressors outside the gene regulatory network can modulate the four fundamental processes. Second, the four proteins interact to form complexes to modulate these processes. Third, other complexation events contribute non-linearly too.

#### S2.2 Modelling protein complexation

Our model describes protein complexation in accordance with the procedures explained in Chapter 7 of [50]. The underlying assumption is that protein-protein interactions occur on a much faster timescale than the functions (transcription and translation) carried out by proteins and protein complexes, as well as their degradation. This leads to the simplifying assumption that protein-protein interactions are in equilibrium on the model’s slower timescale. The equilibrium dissociation constant of each complex describes the equilibrium of the relevant binding event. For example, applying this simplification to complex A, the fraction of unbound MYCN is as follows:

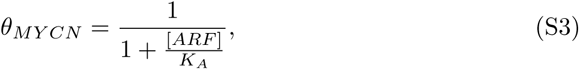

where [*ARF*] is the protein concentration of ARF.

Within our model, MYCN can only form one complex by binding to ARF (Complex A). Other proteins, especially MDM2, can exist in multiple complexes, sometimes with more than one complexation partner. The assumption underlying equation S3 can be generalised to this case, but the unbound fraction of such a protein is expressed in a more complex form. Again, the derivation procedures are explained in Chapter 7 of [50]. For example, the unbound fraction of MDM2 is as follows:

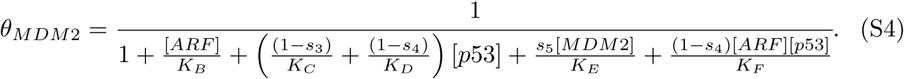

Due to its ability to reside in multiple complexes, MDM2 cannot be fully understood in terms of a binary status (bound or unbound). The bound fraction of MDM2 is divided into further fractions. For example, the fraction of MDM2 in the dimer complex B is 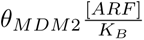. Complex F is a trimer, so the associated fraction is more complex: 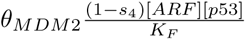. Again, the derivation procedures are explained in Chapter 7 of [50].

Equation S4 also exemplifies how five of the six environmental stressors (all except *s*_1_) are represented in the model. As reviewed, *s*_3_ prevents the formation of complex C, *s*_4_ prevents the formation of complex D, and *s*_5_ promotes the formation of complex E. Promotion and prevention are modelled by multiplying a term by the associated stressor variable (such as *s*_5_) and its complement (such as 1 − *s*_3_) respectively. Conceptually, these environmental stressors are like switches. They modulate some of the cellular processes described by the model in an abstract manner, but they do not represent specific molecular species.

Derived similarly, the following equations are the fractions of unbound p53 and ARF:

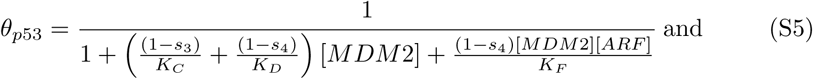

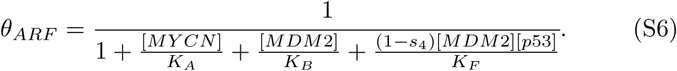

#### S2.3 Modelling protein-DNA interactions

Mathematically, these binding events are handled similarly to protein complexation, but the equilibrium dissociation constants are labelled numerically rather than alphabetically.

Cooperative binding is a mechanism that regulates protein-DNA interactions but not protein complexation. It is modelled by the Hill equation, which is parameterised by the Hill coefficient. When the Hill coefficient is bigger than one, it means that a copy of macromolecule A binds to a copy of macromolecule B more easily when the latter is already bound to other copies of A. Conversely, when the Hill coefficient is smaller than one, a copy of B loses some of its affinity for A with each binding event. When it is exactly one, cooperative behaviour is irrelevant. Most of the binding events described by our model are assumed to be non-cooperative, so they are represented by Michaelis–Menten terms. Examples are equations S3 and S4.

One major exception is the interaction between p53 and the genome. When p53 binds to DNA, it does so sequence-specifically as a tetramer and occupies two consecutive half-sites [117, 118]. The Hill coefficient for this interaction is 1.8 [118, 119]. We will illustrate how the Hill equation is incorporated into the terms associated with p53-DNA binding with an example. The promoter region of *p53* is either free (unbound), bound to p53, bound to MYCN, or bound to both proteins. It is assumed in our model that when there are two transcriptional activators for a promoter, they bind to the promoter independently. Therefore, MYCN [25] and p53 [88] do not interact with DNA cooperatively in our model. The following equation models the fraction of *p53* promoters that are unbound:

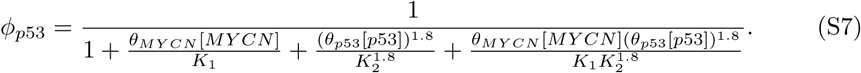

Another exception is the influence of *s*_1_ on the promoter region of *ARF*. In a stressed cell, *ARF* transcription is upregulated by proteins such as E2F-1, *β*-catenin, and Ras [60, 64, 65]. Earlier in subsection S2.2, we illustrated how five of the six environmental stressors are represented as simple switches in our model, that they do not represent specific molecular species. The stressor *s*_1_, which upregulates *ARF* transcription, is an exception. Within our model, it acts like an abstract transcription factor representing the set of proteins that upregulate *ARF* transcription. It can interact with the other molecular species in the model, but there is no evidence of cooperative binding in these interactions, so Michaelis–Menten kinetics is assumed. The fraction of *ARF* promoters that are unbound is given by the following equation:

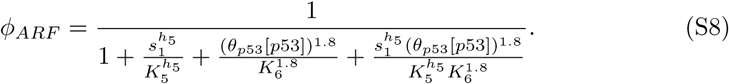

Derived similarly, the following equation is the fraction of *MDM2* promoters that are unbound:

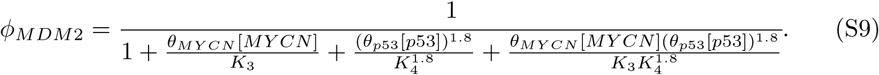

#### S2.4 Modelling protein-mRNA interactions

The fraction of unbound *MYCN* mRNA is given by the following equation:

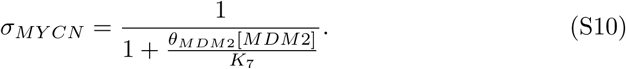

Although the positive interaction between MDM2 and *p53* mRNA can be described by a Michaelis–Menten term as usual, the negative and indirect influence of MDM2 on *p53* translation must be treated abstractly because L26 is not in the model. The solution is to assume that MDM2 either ‘occupies’ an activating site or an inhibiting site on a *p53* mRNA transcript depending on whether the cell is stressed or not. As explained in subsection S2.2, *s*_2_ acts like a simple switch modulating the two sites’ competition for MDM2. This representation is not mechanistically accurate, but it captures the functional consequences of MDM2’s dual role in a phenomenological sense (different inputs give different outputs).

Based on these principles, the following equation gives us the fraction of unbound *p53* mRNA:

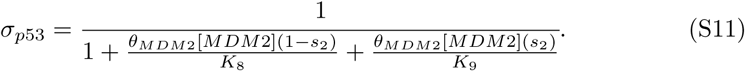

#### S2.5 Modelling functional effects of complexation

Our model describes the functional effects of complexation using intuitive additive terms. When it is in a complex, a protein may degrade at a different rate or be incapable of transcribing another gene (transactivation). For example, MYCN degrades at two different rates, *d_MY CN_* (fast) and *d_MY CN−A_* (slow), in its unbound and bound states respectively. Therefore, the overall protein degradation rate of MYCN is [*θ_MY CN_ d_MY CN_* + (1 − *θ_MY CN_*)*d_MY CN −A_*][*MY CN*].

#### S2.6 Model parameters

The model parameters can be divided into two major categories. The first category contains the rates of mRNA production, mRNA degradation, protein synthesis, and protein degradation in equations 1 to 8. The second category contains the equilibrium dissociation constants parameterising the myriad of complexation events described in subsections 2.1.2, 2.1.3, and S2.4 in the main manuscript and subsections S2.2, S2.3, and S2.4 in the supplementary file.

##### S2.6.1 Fundamental processes

The natural mRNA and protein degradation rates, denoted by *d_m_* and *d_p_*, are 0.06 h^-1^ and 0.05 h^-1^ respectively [51]. It is assumed that *MYCN* mRNA transcripts degrade 10 times more slowly than the natural rate (*d_MY CNm_* = 0.1*d_m_* = 0.006 h^-1^). In complex C, p53 degrades faster than usual. In complexes B and E, MDM2 degrades faster than usual. It is assumed that these three enhanced protein degradation rates are 10 times higher than the natural rate (*d_p_*_53*−*_*_C_* = *d_MDM_*_2*−*_*_BE_* = 10*d_p_* = 0.5 h^-1^).

By setting the derivatives in equations S1 and S2 to zero, it is possible to express the steady-state mRNA and protein abundance levels as 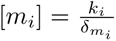 and 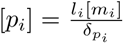 respectively. In order to estimate the transcription and translation rates of each species from the two expressions, we need the steady-state values of [*m_i_*] and [*p_i_*]. The Gene Expression Omnibus (GEO) repository contains mRNA sequencing [52] and ribosome profiling [53] datasets. Specifically, one dataset (accession number: GSE89413) records mRNA abundance levels in NB cells taken from different cell lines (including SH-SY5Y) as reads *per* kilobase million (RPKM), whilst a ribosome profiling dataset (accession number: GSE119615) records protein abundance levels in NB cells taken from the SH-SY5Y cell line as RPKM.

First, we discuss the transcription rates. The steady-state mRNA abundance level ([*m_i_*]) of gene *i* is proportional to its RPKM value (*r_i_*) in the mRNA sequencing dataset divided by the sum of all the RPKM values (Σ*r_i_*) therein. This relative transcription level can be made absolute by multiplication with the total copy number of mRNA in a cell (*N_m_* = 2.3 × 10^4^ copies *per* cell [51]). Therefore, [*m_i_*] = *r_i_N_m_/Σr_i_*. This quantity is multiplied by the natural mRNA degradation rate (*δ_mi_* = *d_m_* = 0.06 h^-1^) to obtained *k_i_* = [*m_i_*]*δ_m_*.

Considering the mRNA sequencing dataset describes the overall behaviour of NB cells, the model assumes that each estimate given by *k_i_* = [*m_i_*]*δ_mi_* corresponds to the highest transcription rate available to a gene when there are more than one rate. The estimates are denoted by *α_MYCN_*, *β_p53_*, *β_MDM2_*, and *β_ARF_*, which represent the natural transcription rate of *MYCN* and the enhanced transcription rates of the other three genes. In our model, *MYCN* is always transcribed naturally and the other three genes are transcribed faster when their activating mechanisms are active. In addition to having a fixed transcription rate, *MYCN* is a special case because the copy number of *MYCN* (*g_MY CN_*) is hidden in the estimate *k_MY CN_*, meaning *k_MY CN_* = *g_MY CN_ α_MY CN_*. The SH-SY5Y cell line is not MA [120], so *g_MY CN_* = 1 and *k_MY CN_* = *α_MY CN_* for the purpose of parameterisation. The natural transcription rates of *p53*, *MDM2*, and *ARF* (*α_p53_*, *α_MDM2_*, and *α_ARF_*) are assumed to be 10 times slower than the corresponding enhanced rates.

Moving on to the translation rates, the RPKM values ([*p_i_*]) in the ribosome profiling dataset can be used in a similar manner. First, [*p_i_*]*/Σ*[*p_i_*] estimates the relative translation level of gene *i*. Multiplying this estimate by the total number of proteins in a cell (*N_p_* = 7.5 × 10^9^ proteins *per* cell [51]) makes it absolute. Multiplying the absolute translation level by *δ_pi_ /*[*m_i_*] gives the translation rate of gene *i* as *l_i_* = (*δ_pi_ /*[*m_i_*])(*N_p_*[*p_i_*]*/Σ*[*p_i_*]). The appearance of [*m_i_*] in the denominator means the rate is independent of the amount of mRNA. The natural protein degradation rate is *d_p_* = 0.05 h^-1^. The steady-state abundance level of the corresponding mRNA [*m_i_*] is as described in the previous paragraph.

Considering the dataset describes the overall behaviour of NB cells, the model assumes that each estimate *l_i_* = (*δ_pi_ /*[*m_i_*])(*N_p_*[*p_i_*]*/Σ*[*p_i_*]) corresponds to the highest translation rate available to a gene. Therefore, the four estimates are *λ_MYCN_*, *µ_p53_*, *λ_MDM2_*, and *λ_ARF_*. Apart from p53, each of them is produced at just one natural rate. p53 has a natural translation rate (*λ_p53_*), a reduced translation rate (*κ_p53_*), and an enhanced translation rate (*µ_p53_*). The estimate is *µ_p53_* and the natural rate *λ_p53_* and reduced rate *κ_p53_* are assumed to be 10 and 100 times slower respectively.

##### S2.6.2 Equilibrium dissociation constants and Hill coefficient

As reported in table 2, literature values are available for many of these parameters. Two exceptions are *K_A_* and *K_B_*, so the model assumes a value of 100 nM based on the average order of magnitude of the other equilibrium dissociation constants. *K*_5_ is the equilibrium dissociation constant for the interaction between the abstract transcription factor *s*_1_; representing E2F-1, *β*-catenin, and Ras; and the *ARF* gene’s promoter region. Since *s*_1_ is dimensionless and ranges from zero to one, *K*_5_ = 1.

#### S2.7 Computer simulations

We conducted three sets of computer simulations. First, we varied three groups of parameters with and without *MYCN* amplification in a grid search. The results were used to confirm the model’s stable long-term behaviour (equilibrium), which represents the clinical/treatment outcome; understand the outcome’s sensitivity to the three groups of parameters; and establish a NB classification system based on our hypothesis. Second, we focused on the two most sensitive groups of parameters, repeating the grid search without *MYCN* amplification and at five levels of *MYCN* amplification. Third, we validated the conclusions of the first two experiments by exploring the parametric space randomly.

##### S2.7.1 First grid search

In the first experiment, we varied three groups of parameters together and for each parametric combination, we tested the model’s long-term stability.

- We tested 2^6^ = 64 input combinations after binarising the six stressors so that *s_i_* = 0.9 or *s_i_* = 0.
- The unknown Hill coefficient *h*_5_ was strategically varied from 0.2 (negative cooperativity) to one (no cooperativity) and then to four (positive cooperativity) to cover all three regimes.
- We tested three values for each of the four transcription rates (*α_MYCN_*, *β_p53_*, *β_MDM2_*, and *β_ARF_*) and four basal translation rates (*λ_MYCN_*, *µ_p53_*, *λ_MDM2_*, and *λ_ARF_*) based on experimental measurements. For each rate, the estimate in table 1, 70 % of the estimate, and 130 % of the estimate were tested. For a process with two or three possible rates, all of them were varied to maintain the ratios between them. For example, when 70 % of the estimate of *µ_p53_* was used, *λ_p53_* was also reduced by 30 %. Within group three, we tested 3^8^ = 6561 combinations of rates.

The equilibrium dissociation constants were fixed under the assumption that they do not vary between two patients. Since *s*_1_ is relevant to *h*_5_ in the model, when *h*_5_ was varied (0.2, one, and four) with the other stressors off (zero) and the rates at their default values, *s*_1_ was set to an additional value: 0.5. It means three additional parametric combinations were tested. In total, 64 · 3 · 6561 + 3 = 1259715 parametric combinations were tested in the first experiment.

Each parametric combination was used to perform two sets of 10 simulations within the MATLAB environment with the ode45 solver function, once with *g_MYCN_* = 1 and once with *g_MYCN_* = 2. For each combination and each *MYCN* amplification status, 10 simulations were conducted from 10 random initial conditions. An initial condition was created by sampling each of the eight variables randomly from the continuous uniform distribution in the range [0, 1]. A simulation was terminated at *t* = 10^6^ or when an equilibrium was reached before *t* = 10^6^. Equilibria were detected by computing the ratio between the norm of the state variables’ derivatives and the norm of the state variables themselves at the end of each time step during the simulations. When the ratio fell below the cutoff threshold of 10*^−^*^6^, the simulation was terminated. In mathematical terms, these equilibria are called stable fixed points, which correspond to clinically observable and relevant scenarios.

The identified fixed points’ stability was tested numerically. Starting from each fixed point, we perturbed each state variable to a random value in the range between zero and 10 times the equilibrium value. The simulation was repeated from this perturbed condition and if the system (eight-dimensional vector comprising the state variables) returned to the fixed point, the fixed point was classified as stable.

##### S2.7.2 Second grid search

In the second experiment, we wanted to learn more about how the three groups of parameters interact to affect the system’s long-term behaviour. Informed by the results of the first experiment, we set *h*_5_ = 1 and varied *s_i_* and the rates together. We did so at six levels of *MYCN* amplification, including the wild-type genotype. Within the MATLAB environment, 64 · 6561 = 419904 combinations of the parametric values tested in the first experiment were used to perform six sets of simulations with the ode45 solver function, one for each of *g_MYCN_* = 1, *g_MYCN_* = 2, *g_MYCN_* = 3, *g_MYCN_* = 4, *g_MYCN_* = 6, and *g_MYCN_* = 8. The same initialisation and termination procedures were adhered to.

##### S2.7.3 Random exploration of the parametric space

In the third experiment, in order to validate the conclusions of the two grid search experiments, we generalised them by sampling from the whole parametric space. We did so by varying *s_i_* and the rates together and randomly, whilst keeping *h*_5_ = 1, leading to 10000 random parametric combinations. Each combination was generated by sampling each stressor and rate randomly and uniformly between the minimum and maximum values tested in the first two experiments. Then, like the second experiment, each parametric combination was used to perform six simulations with *g_MYCN_* = 1, *g_MYCN_* = 2, *g_MYCN_* = 3, *g_MYCN_* = 4, *g_MYCN_* = 6, and *g_MYCN_* = 8. The other procedures followed in the second experiment were adhered to too.

#### S2.8 Data mining and visualisation

Experiment two. The Apriori algorithm computes three metrics in order to identify frequent itemsets and association rules in each subset of parametric combinations: support, confidence, and lift. We will illustrate them using the parameters *s*_1_ and *s*_2_ in the subset of WT equilibria predicted to have good clinical/treatment outcomes.

- The support of *s*_1_ being on (0.9) and *s*_2_ being off (0) in this subset is the number of parametric combinations where *s*_1_ = 0.9 and *s*_2_ = 0 divided by the total number of combinations in this subset. If the support is above a user-defined threshold, the combination of *s*_1_ being on (0.9) and *s*_2_ being off (0) is a frequent itemset.
- The confidence of *s*_1_ being on (0.9) with respect to *s*_2_ being off (0) is the number of combinations where *s*_1_ = 0.9 and *s*_2_ = 0 divided by the number of combinations where *s*_2_ = 0 in the subset. In other words, it is the support of *s*_1_ being on (0.9) and *s*_2_ being off (0) divided by the support of *s*_2_ being off (0). This metric tells you how likely it is that *s*_1_ = 0.9 in a parametric combination if *s*_2_ = 0 therein in this subset. A major problem with this metric is that it is sensitive to the frequency of an item in a dataset. For example, if *s*_1_ = 0.9 throughout the subset, the confidence of *s*_1_ being on (0.9) with respect to *s*_2_ being off (0) will of course be 100 %, but it does not buttress the claim that the likelihood of the former is boosted by the truth of the latter.
- The lift mitigates this problem of the confidence metric by taking the baseline occurrence of the item being boosted into account. For example, the lift of *s*_1_ being on (0.9) with respect to *s*_2_ being off (0) takes the baseline occurrence of *s*_1_ being on (0.9) in the subset into account. It is the confidence of *s*_1_ being on (0.9) with respect to *s*_2_ being off (0) divided by the support of *s*_1_ being on (0.9). If the lift is above one, it means that *s*_2_ being off (0) increases the likelihood of *s*_1_ being on (0.9) in a parametric combination in the subset. This association rule is denoted in this way: *s*_2_ = 0 ⇒ *s*_1_ = 0.9.

Experiment three. Using the ggpairs function within the GGally R package, we created an augmented scatterplot matrix (also called a ggplot2 generalised pairs plot) for each subset of random parametric combinations. It is a 14×14 matrix because in the third experiment, we took random samples from a 14-dimensional parametric space (six stressors and eight rates). Below the leading diagonal are the 91 scatterplots comparing the unique pairs formed by the 14 parameters. On the diagonal are the probability density functions of the 14 parameters in this random sample. Above the diagonal, the Pearson correlation coefficients of the 91 pairs of parameters are reported. Each correlation’s statistical significance is shown too. In a matrix element, *** means the reported p-value is smaller than 0.001; ** means that it is smaller than 0.01; * means that it is smaller than 0.05; and a full stop (.) means it is smaller than 0.10.

### S3 Additional results

Below are the four sets of 20 association rules (subsection 3.4) extracted from the results of the second experiment with the Apriori algorithm. We divided the fixed points found in the experiment into four subsets (genotype-outcome combinations): WT (*g_MYCN_* = 1) or MA (*g_MYCN_* = 2) and good (p53/MYCN above 3.5) or poor (p53/MYCN below 3.5) clinical/treatment outcomes. We applied the Apriori algorithm to the parametric combinations that gave us the fixed points in each subset.

**Table S1:**
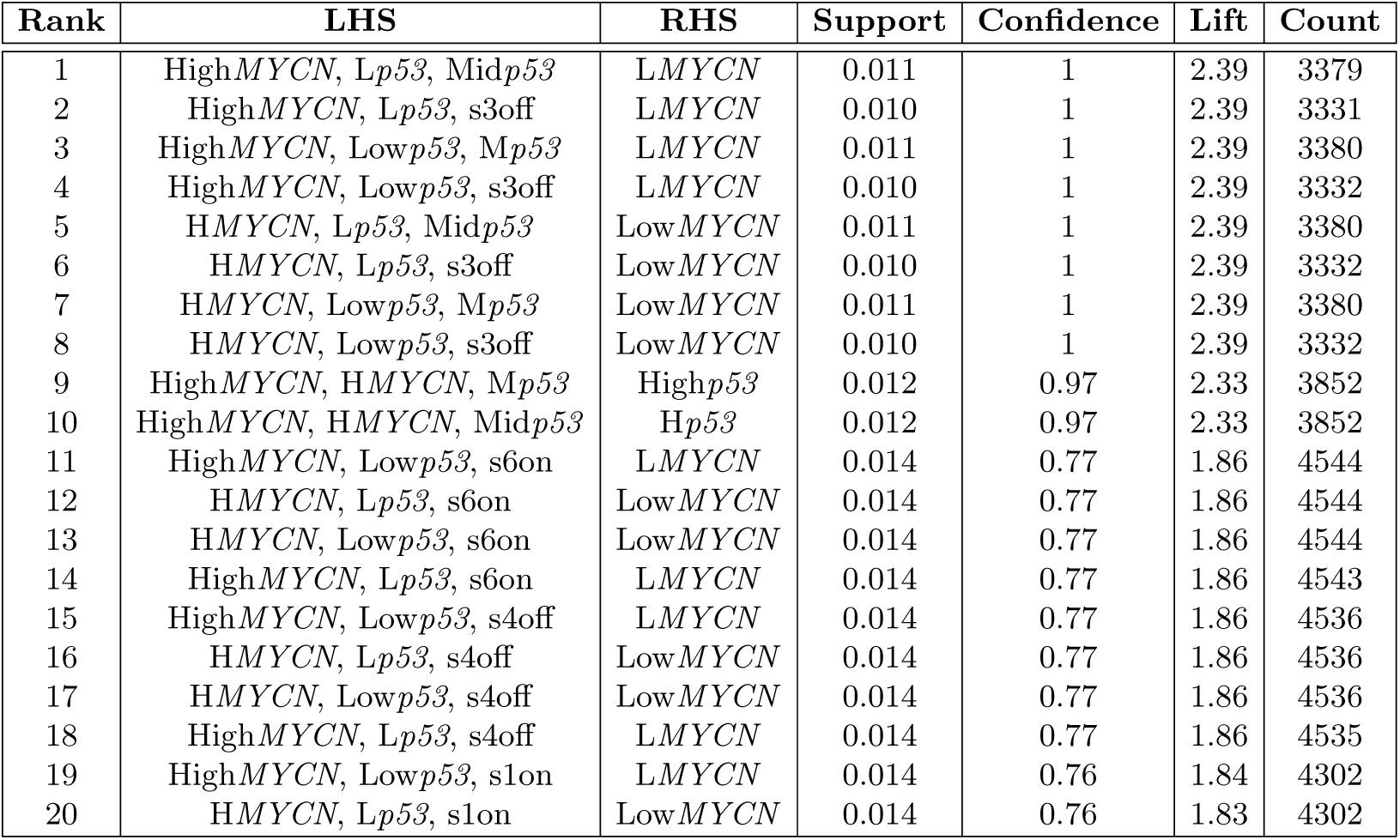
Top 20 association rules regarding the parametric combinations associated with WT fixed points with good outcomes in the second experiment. L*X*, M*X*, H*X* represent transcription rates of *X* that are 70%, 100%, 130% of the default (also called middle) value. Low*X*, Mid*X*, High*X* represent translation rates of *X* that are 70%, 100%, 130% of the default (also called middle) value. sion and sioff represent the active and inactive statuses of *s_i_* respectively. LHS denotes the antecedent (left-hand side) of a rule, whilst RHS denotes the consequent (right-hand side) of a rule.

| Rank | LHS | RHS | Support | Confidence | Lift | Count |
| --- | --- | --- | --- | --- | --- | --- |
| 1 | HighMYCN, Lp53, Midp53 | LMYCN | 0.011 | 1 | 2.39 | 3379 |
| 2 | HighMYCN, Lp53, s3off | LMYCN | 0.010 | 1 | 2.39 | 3331 |
| 3 | HighMYCN, Lowp53, Mp53 | LMYCN | 0.011 | 1 | 2.39 | 3380 |
| 4 | HighMYCN, Lowp53, s3off | LMYCN | 0.010 | 1 | 2.39 | 3332 |
| 5 | HMYCN, Lp53, Midp53 | LowMYCN | 0.011 | 1 | 2.39 | 3380 |
| 6 | HMYCN, Lp53, s3off | LowMYCN | 0.010 | 1 | 2.39 | 3332 |
| 7 | HMYCN, Lowp53, Mp53 | LowMYCN | 0.011 | 1 | 2.39 | 3380 |
| 8 | HMYCN, Lowp53, s3off | LowMYCN | 0.010 | 1 | 2.39 | 3332 |
| 9 | HighMYCN, HMYCN, Mp53 | Highp53 | 0.012 | 0.97 | 2.33 | 3852 |
| 10 | HighMYCN, HMYCN, Midp53 | Hp53 | 0.012 | 0.97 | 2.33 | 3852 |
| 11 | HighMYCN, Lowp53, s6on | LMYCN | 0.014 | 0.77 | 1.86 | 4544 |
| 12 | HMYCN, Lp53, s6on | LowMYCN | 0.014 | 0.77 | 1.86 | 4544 |
| 13 | HMYCN, Lowp53, s6on | LowMYCN | 0.014 | 0.77 | 1.86 | 4544 |
| 14 | HighMYCN, Lp53, s6on | LMYCN | 0.014 | 0.77 | 1.86 | 4543 |
| 15 | HighMYCN, Lowp53, s4off | LMYCN | 0.014 | 0.77 | 1.86 | 4536 |
| 16 | HMYCN, Lp53, s4off | LowMYCN | 0.014 | 0.77 | 1.86 | 4536 |
| 17 | HMYCN, Lowp53, s4off | LowMYCN | 0.014 | 0.77 | 1.86 | 4536 |
| 18 | HighMYCN, Lp53, s4off | LMYCN | 0.014 | 0.77 | 1.86 | 4535 |
| 19 | HighMYCN, Lowp53, s1on | LMYCN | 0.014 | 0.76 | 1.84 | 4302 |
| 20 | HMYCN, Lp53, s1on | LowMYCN | 0.014 | 0.76 | 1.83 | 4302 |

**Table S2:** Top 20 association rules regarding the parametric combinations associated with WT fixed points with poor outcomes in the second experiment. L*X*, M*X*, H*X* represent transcription rates of *X* that are 70%, 100%, 130% of the default (also called middle) value. Low*X*, Mid*X*, High*X* represent translation rates of *X* that are 70%, 100%, 130% of the default (also called middle) value. sion and sioff represent the active and inactive statuses of *s_i_* respectively. LHS denotes the antecedent (left-hand side) of a rule, whilst RHS denotes the consequent (right-hand side) of a rule.

| Rank | LHS | RHS | Support | Confidence | Lift | Count |
| --- | --- | --- | --- | --- | --- | --- |
| 1 | Hp53, s3on | Lowp53 | 0.076 | 1 | 2.16 | 2592 |
| 2 | Highp53, s3on | Lp53 | 0.0769 | 1 | 2.16 | 2592 |
| 3 | Hp53, s3on, s6on | Lowp53 | 0.038 | 1 | 2.16 | 1296 |
| 4 | Hp53, s3on, s4off | Lowp53 | 0.038 | 1 | 2.16 | 1296 |
| 5 | Hp53, s2on, s3on | Lowp53 | 0.038 | 1 | 2.16 | 1296 |
| 6 | Hp53, s1on, s3on | Lowp53 | 0.038 | 1 | 2.16 | 1296 |
| 7 | Hp53, s3on, s5off | Lowp53 | 0.038 | 1 | 2.16 | 1296 |
| 8 | Hp53, s3on, s5on | Lowp53 | 0.0387 | 1 | 2.16 | 1296 |
| 9 | Hp53, s1off, s3on | Lowp53 | 0.038 | 1 | 2.16 | 1296 |
| 10 | Hp53, s2off, s3on | Lowp53 | 0.038 | 1 | 2.16 | 1296 |
| 11 | Hp53, s3on, s4on | Lowp53 | 0.038 | 1 | 2.16 | 1296 |
| 12 | Hp53, s3on, s6off | Lowp53 | 0.038 | 1 | 2.16 | 1296 |
| 13 | HMYCN, Hp53, s3on | Lowp53 | 0.076 | 1 | 2.16 | 2592 |
| 14 | HighMYCN, Hp53, s3on | Lowp53 | 0.076 | 1 | 2.16 | 2592 |
| 15 | Highp53, s3on, s6on | Lp53 | 0.038 | 1 | 2.16 | 1296 |
| 16 | Highp53, s3on, s4off | Lp53 | 0.038 | 1 | 2.16 | 1296 |
| 17 | Highp53, s2on, s3on | Lp53 | 0.0384 | 1 | 2.16 | 1296 |
| 18 | Highp53, s1on, s3on | Lp53 | 0.0384 | 1 | 2.16 | 1296 |
| 19 | Highp53, s3on, s5off | Lp53 | 0.038 | 1 | 2.16 | 1296 |
| 20 | Highp53, s3on, s5on | Lp53 | 0.038 | 1 | 2.16 | 1296 |

**Table S3:** Top 20 association rules regarding the parametric combinations associated with MA fixed points with good outcomes in the second experiment. L*X*, M*X*, H*X* represent transcription rates of *X* that are 70%, 100%, 130% of the default (also called middle) value. Low*X*, Mid*X*, High*X* represent translation rates of *X* that are 70%, 100%, 130% of the default (also called middle) value. sion and sioff represent the active and inactive statuses of *s_i_* respectively. LHS denotes the antecedent (left-hand side) of a rule, whilst RHS denotes the consequent (right-hand side) of a rule.

| Rank | LHS | RHS | Support | Confidence | Lift | Count |
| --- | --- | --- | --- | --- | --- | --- |
| 1 | HighMYCN, Midp53 | Hp53 | 0.023 | 1 | 1.85 | 2920 |
| 2 | HMYCN, Midp53 | Hp53 | 0.023 | 1 | 1.85 | 2920 |
| 3 | HighMYCN, Midp53, s3on | Hp53 | 0.021 | 1 | 1.85 | 2612 |
| 4 | HighMYCN, LMYCN, Midp53 | Hp53 | 0.023 | 1 | 1.85 | 2900 |
| 5 | HMYCN, Midp53, s3on | Hp53 | 0.021 | 1 | 1.85 | 2612 |
| 6 | HMYCN, LowMYCN, Midp53 | Hp53 | 0.023 | 1 | 1.85 | 2900 |
| 7 | HighMYCN, LMYCN, Midp53, s3on | Hp53 | 0.021 | 1 | 1.85 | 2592 |
| 8 | HMYCN, LowMYCN, Midp53, s3on | Hp53 | 0.021 | 1 | 1.85 | 2592 |
| 9 | HighMYCN, Mp53 | Highp53 | 0.023 | 1 | 1.85 | 2920 |
| 10 | HMYCN, Mp53 | Highp53 | 0.023 | 1 | 1.85 | 2920 |
| 11 | HighMYCN, Mp53, s3on | Highp53 | 0.021 | 1 | 1.85 | 2612 |
| 12 | HighMYCN, LMYCN, Mp53 | Highp53 | 0.023 | 1 | 1.85 | 2900 |
| 13 | HMYCN, Mp53, s3on | Highp53 | 0.021 | 1 | 1.85 | 2612 |
| 14 | HMYCN, LowMYCN, Mp53 | Highp53 | 0.023 | 1 | 1.85 | 2900 |
| 15 | HighMYCN, LMYCN, Mp53, s3on | Highp53 | 0.021 | 1 | 1.85 | 2592 |
| 16 | HMYCN, LowMYCN, Mp53, s3on | Highp53 | 0.021 | 1 | 1.85 | 2592 |
| 17 | Lowp53, MidMYCN | Hp53 | 0.010 | 1 | 1.85 | 1254 |
| 18 | Lowp53, MMYCN | Hp53 | 0.010 | 1 | 1.85 | 1255 |
| 19 | HighMYCN, Midp53, s4off | Hp53 | 0.011 | 1 | 1.85 | 1460 |
| 20 | HighMYCN, Midp53, s6on | Hp53 | 0.011 | 1 | 1.85 | 1466 |

**Table S4:**
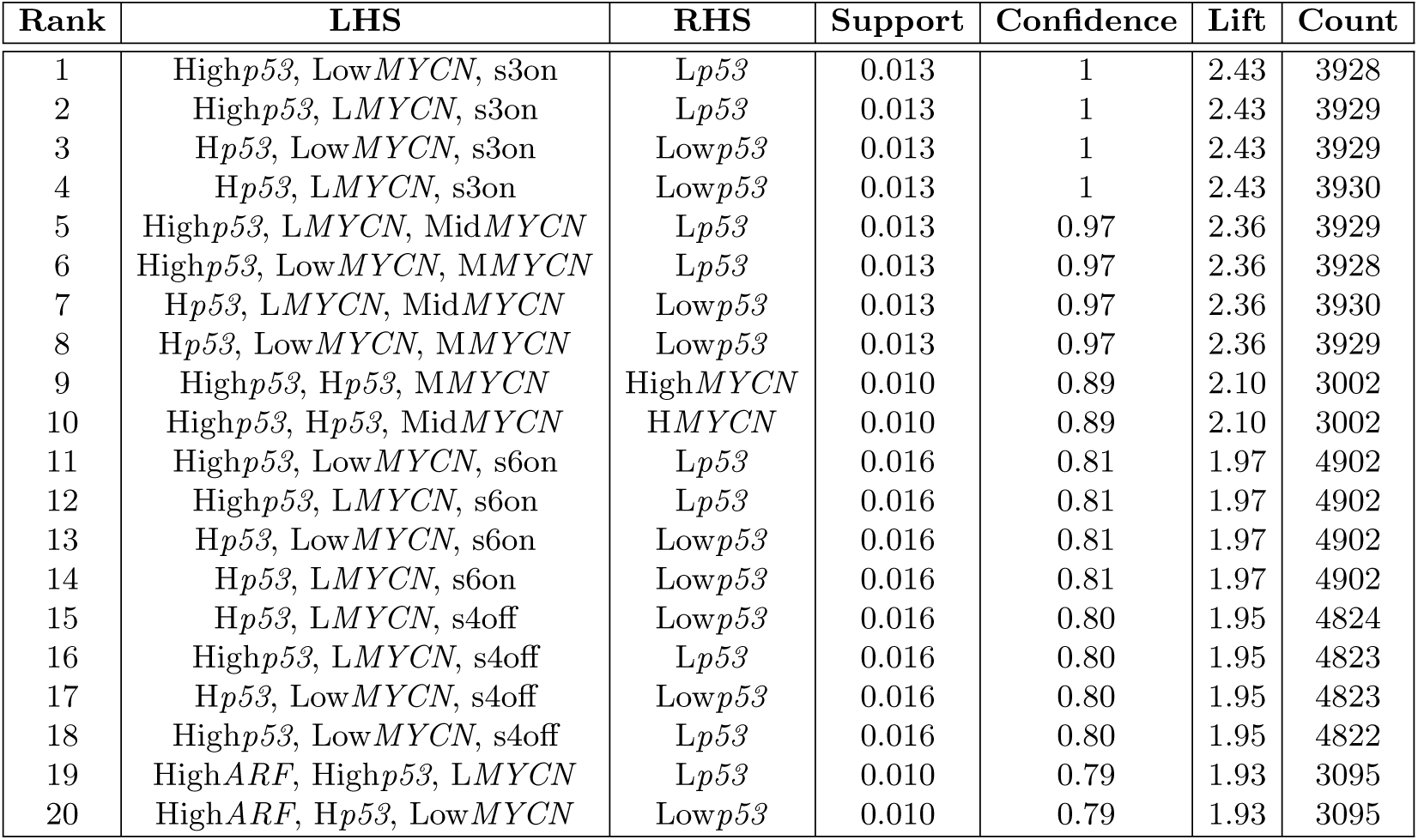
Top 20 association rules regarding the parametric combinations associated with MA fixed points with poor outcomes in the second experiment. L*X*, M*X*, H*X* represent transcription rates of *X* that are 70%, 100%, 130% of the default (also called middle) value. Low*X*, Mid*X*, High*X* represent translation rates of *X* that are 70%, 100%, 130% of the default (also called middle) value. sion and sioff represent the active and inactive statuses of *s_i_* respectively. LHS denotes the antecedent (left-hand side) of a rule, whilst RHS denotes the consequent (right-hand side) of a rule.

